# Single-cell analysis of human diversity in circulating immune cells

**DOI:** 10.1101/2024.06.30.601119

**Authors:** Kian Hong Kock, Le Min Tan, Kyung Yeon Han, Yoshinari Ando, Damita Jevapatarakul, Ankita Chatterjee, Quy Xiao Xuan Lin, Eliora Violain Buyamin, Radhika Sonthalia, Deepa Rajagopalan, Yoshihiko Tomofuji, Shvetha Sankaran, Mi-So Park, Mai Abe, Juthamard Chantaraamporn, Seiko Furukawa, Supratim Ghosh, Gyo Inoue, Miki Kojima, Tsukasa Kouno, Jinyeong Lim, Keiko Myouzen, Sarintip Nguantad, Jin-Mi Oh, Nirmala Arul Rayan, Sumanta Sarkar, Akari Suzuki, Narita Thungsatianpun, Prasanna Nori Venkatesh, Jonathan Moody, Masahiro Nakano, Ziyue Chen, Chi Tian, Yuntian Zhang, Yihan Tong, Crystal T.Y. Tan, Anteneh Mehari Tizazu, SG10K_Health Consortium, Marie Loh, You Yi Hwang, Roger C Ho, Anis Larbi, Tze Pin Ng, Hong-Hee Won, Fred A. Wright, Alexandra-Chloé Villani, Jong-Eun Park, Murim Choi, Boxiang Liu, Arindam Maitra, Manop Pithukpakorn, Bhoom Suktitipat, Kazuyoshi Ishigaki, Yukinori Okada, Kazuhiko Yamamoto, Piero Carninci, John C. Chambers, Chung-Chau Hon, Ponpan Matangkasombut, Varodom Charoensawan, Partha P. Majumder, Jay W. Shin, Woong-Yang Park, Shyam Prabhakar

**Author notes:** Corresponding authors. Correspondence to <u>Shyam Prabhakar</u>, <u>Woong-Yang Park</u>, <u>Jay W.</u> <u>Shin</u>. These authors contributed equally (Le Min Tan, Kyung Yeon Han, Yoshinari Ando, Damita Jevapatarakul, Ankita Chatterjee, Quy Xiao Xuan Lin). These authors contributed equally (Eliora Violain Buyamin, Radhika Sonthalia, Deepa Rajagopalan, Yoshihiko Tomofuji).

## Abstract

Lack of diversity and proportionate representation in genomics datasets and databases contributes to inequity in healthcare outcomes globally^1,2^. The relationships of human diversity with biological and biomedical phenotypes are pervasive^3^, yet remain understudied, particularly in a single-cell genomics context. Here we present the Asian Immune Diversity Atlas (AIDA), a multi-national single-cell RNA-sequencing (scRNA-seq) healthy reference atlas of human immune cells. AIDA comprises 1,265,624 circulating immune cells from 619 healthy donors and 6 controls, spanning 7 population groups across 5 countries. AIDA is one of the largest healthy blood datasets in terms of number of cells, and also the most diverse in terms of number of population groups. Though population groups are frequently compared at the continental level, we identified a pervasive impact of sub-continental diversity on cellular and molecular properties of immune cells. These included cell populations and genes implicated in disease risk and pathogenesis as well as those relevant for diagnostics. We detected single-cell signatures of human diversity not apparent at the level of cell types, as well as modulation of the effects of age and sex by self-reported ethnicity. We discovered functional genetic variants influencing cell type-specific gene expression, including context-dependent effects, which were under-represented in analyses of non-Asian population groups, and which helped contextualise disease-associated variants. We validated our findings using multiple independent datasets and cohorts. AIDA provides fundamental insights into the relationships of human diversity with immune cell phenotypes, enables analyses of multi-ancestry disease datasets, and facilitates the development of precision medicine efforts in Asia and beyond.

## Introduction

Humans are diverse in all respects. Our molecular diversity drives differences in our cellular traits, which in turn feeds into differences in how our bodies develop, function, and respond to disease. Molecular variation across individuals is not random; rather, it correlates with ancestry, age, genetics, sex, environment, and lifestyle^1^, though in ways we do not fully understand. One consequence is that molecular diagnostics that work in one population may not be as effective in another^4^. Moreover, disease risk, pathological processes, and drug responses can vary across populations, due to a complex combination of genetic and environmental differences^1–3^. Consequently, an understanding of human molecular and cellular variation is essential not merely for understanding human biology, but also for personalised medical care and equitable outcomes from biomedical research^5^.

The study of diversity in immune cells across humans is of great interest, since blood cell proportions are routinely used for diagnosis, variation in blood traits is associated with disease risk^6^, and immunophenotyping is utilised to monitor diseases such as HIV/AIDS, leukaemias, and lymphomas^7^. Haematological traits vary across ancestries^8–10^, at least partly due to population-specific genetic variants^11^. However, most existing studies provide limited detail on molecular and gene expression profiles, particularly at single-cell resolution.

Recently, single-cell RNA-sequencing (scRNA-seq) studies have examined ancestry-specific immunological traits of US-based populations for lupus and *in vitro* viral infection^12–14^. In addition, cohort-scale scRNA-seq analyses have identified cell type-specific expression quantitative trait loci (eQTL) linked to GWAS variants^15–17^. However, each of these studies focused on a single country and at most two ancestries. More broadly, despite the pressing need, there is a lack of diversity in genomics datasets^1^. For example, individuals of European ancestry, constituting ∼15% of the world’s population, represented ∼86% of the NHGRI-EBI GWAS Catalog in 2021^2^ and ∼85% of the GTEx v8 dataset^18^. To maximise benefit to global communities, it is important to incorporate and characterise human diversity within reference cell atlases and genomics resources.

To address this challenge, we performed scRNA-seq on peripheral blood mononuclear cells (PBMCs) from 619 healthy donors spanning 7 population groups in 5 countries across Asia, a continent inhabited by 60% of the global population^2^. Our Asian Immune Diversity Atlas (AIDA) cohort includes donors from India, Japan, South Korea, and Thailand, as well as Singapore donors of Chinese (SG_Chinese), Malay (SG_Malay), or Indian (SG_Indian) self-reported ethnicities, and thus encompasses a wide range of ancestries^19–24^. The AIDA cohort incorporates a balance of female and male sex and a wide range of adult ages. We characterised the relationships of human diversity with cellular and molecular variation in immune phenotypes, including cell type proportions, cell neighbourhood abundance, and cell type-specific gene expression profiles. Self-reported ethnicity and sex had comparable effects on cell subtype proportions, while the variance explained by age or body mass index (BMI) was typically lower. Moreover, self-reported ethnicity modulated the effects of age and sex on cellular and molecular profiles. Of the variants we identified by cell type-specific eQTL analysis, ∼7% were present at <5% minor allele frequency in the non-Asian 1,000 Genomes super-populations; these included multiple variants colocalising with Asian GWAS loci. Our datasets are available via the Human Cell Atlas (HCA) and Chan Zuckerberg (CZ) CELLxGENE^25^ data portals and have been used for algorithm development in the context of human diversity^26^. Our datasets have also facilitated analyses of biological pathways, such as escape from X-chromosome inactivation (XCI)^27^, and genetic effects on alternative splicing (Tian *et al.*, submitted). Our findings provide fundamental insights into the relationships of age, self-reported ethnicity, sex, and genetic variants with disease-relevant immune phenotypes, and strengthen the scientific case for functional genomics analyses of diverse populations.

## Results

### scRNA-seq atlas of circulating immune cells from diverse population groups

We examined the CZ CELLxGENE Census (version 2023-12-15)^25^, which comprises the largest collection of standardised single-cell data, including healthy reference datasets important for disease comparisons. Across all 26 healthy blood primary datasets (excluding AIDA), 62.4% of cells were annotated as being from European donors, while self-reported ethnicity information was unknown for 32.1% of cells (**Figure 1A**). This is indicative of both the paucity of proportionate representation of the global population in the incipiency of single-cell reference collections, as well as a lack of granularity in examining population groups. We sought to understand the impact of human diversity, focusing on variation across sub-continental population groups, on single-cell genomics profiles of the AIDA cohort. For each of the 619 AIDA donors (**Tables 1,S1**), we performed 5’ scRNA-seq, B-cell receptor sequencing (BCR-seq), T-cell receptor sequencing (TCR-seq), and genotyping (<u>Methods</u>, **Figure 1B,S1A,B**). To minimise technical confounders, we harmonised donor selection criteria, sample processing, and experimental protocols across the 5 study sites (**Figure 1C**), generated single-cell libraries from pooled samples using genetic multiplexing, and adopted a centralised data processing pipeline (**Figure 1B**). Donors spanned an age range of 19 to 77 years (**Table 1**, **Figure 1D**) and were largely balanced in female and male sex (**Table 1**). Principal component analysis (PCA) of donor Illumina GSAv3 genotype data highlighted the diversity of ancestries in the AIDA cohort (**Figure 1E**).

**Figure 1:**
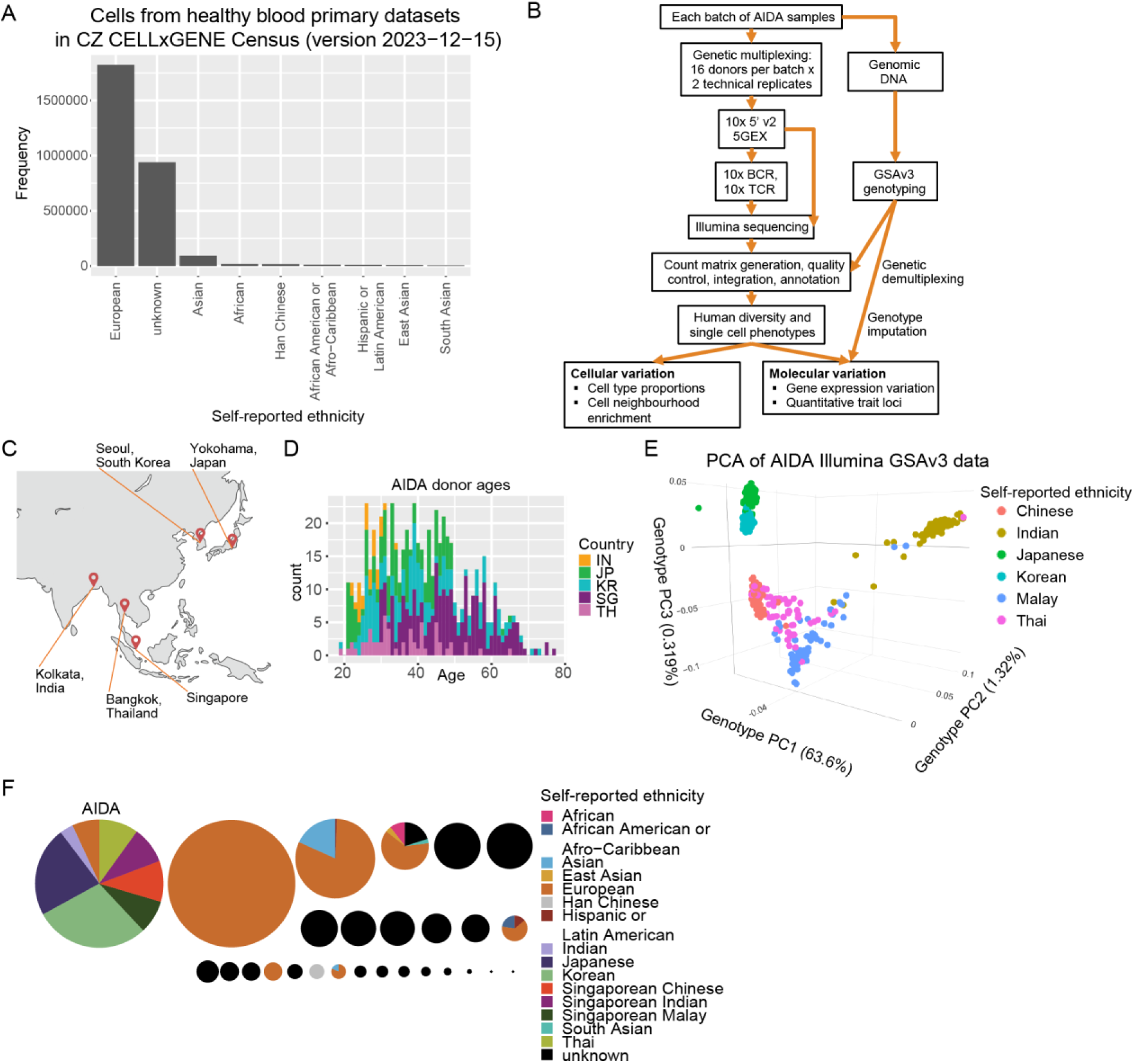
Human diversity in the Asian Immune Diversity Atlas (AIDA) cohort. (**A**) Frequencies of human cells from healthy blood (excluding umbilical cord blood and venous blood) primary datasets in the CZ CELLxGENE Census (version 2023-12-15)^25^, without AIDA, categorised by their self-reported ethnicity metadata. (**B**) AIDA workflow. (**C**) Study site locations. Map adapted from BioRender template (Publication licence agreement number KZ26TPRFSH). (**D**) Histogram of AIDA donor ages, coloured by country (IN:India, JP:Japan, KR:South Korea, SG:Singapore, TH:Thailand). (**E**) Plot of the first three principal components (PCs) from principal component analysis (PCA) of AIDA donor Illumina GSAv3 genotype data, with variance explained by each PC indicated on axis labels. Colours indicate donor self-reported ethnicity. (**F**) Pie charts representing cells in the AIDA dataset, as well as cells from other healthy blood (excluding umbilical cord blood and venous blood) primary datasets in the CZ CELLxGENE Census (version 2023-12-15)^25^. Circle radii are proportional to the numbers of cells in the corresponding datasets; each slice in each pie chart is coloured by the self-reported ethnicity metadata.

**Table 1:** AIDA donor demographics.

| Donor demographics | Number of donors |
| --- | --- |
| Japan | 149 |
| Singaporean Chinese (SG_Chinese) | 85 |
| Singaporean Malay (SG_Malay) | 61 |
| Singaporean Indian (SG_Indian) | 70 |
| South Korea | 165 |
| Thailand | 59 |
| India | 30 |
| **Total** | **619** |
| Female | 348 (56.2%) |
| Male | 271 (43.8%) |
| Age range (years) | 19 to 77 (median=40) |

After doublet removal and cell type-specific quality control (<u>Methods</u>) to remove low-quality cells, we obtained 1,265,624 circulating immune cells. AIDA is one of the largest healthy blood datasets in terms of number of cells, and also the most diverse in terms of number of population groups, relative to existing healthy blood primary datasets in the CZ CELLxGENE Census (version 2023-12-15, **Figure 1F**). We clustered these cells into 8 major immune cell types: B, CD34^+^ haematopoietic stem and progenitor cell (HSPC), myeloid, natural killer (NK), plasma cell, plasmacytoid dendritic cell (pDC), platelet, and T (**Figure 2A**). The distributions of cells in batch-corrected^28^ gene expression space were broadly consistent across all study sites (**Figure S1C**).

**Figure 2:**
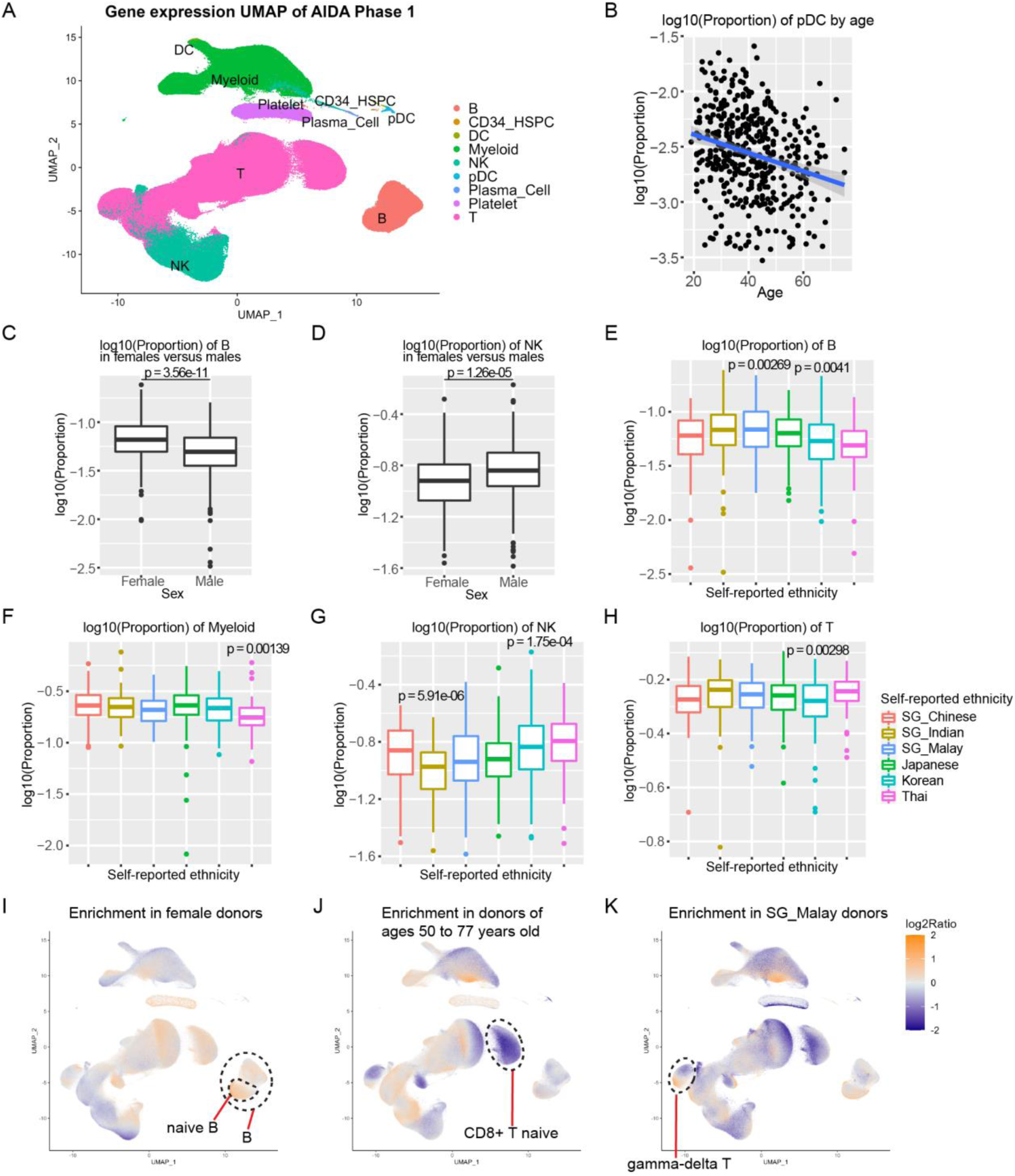
The impact of age, sex, and self-reported ethnicity on AIDA PBMCs. (**A**) Gene expression UMAP of AIDA Data Freeze v2: 1,265,624 PBMCs labelled by major cell type. Data integration was performed using scRNA-seq library as the batch label. (**B**) Scatterplot of plasmacytoid dendritic cell (pDC) proportions against donor age. Linear regression line in blue; grey band indicates the 95% confidence interval. Boxplots depicting distributions of cell type proportions of (**C**) B and (**D**) NK cells in females versus males, relative to all PBMCs per donor. Two-tailed t-test p-values are indicated for the sex covariate in a model of log_10_(Proportion)∼Age+Sex+Self_reported_ethnicity. Boxplots depicting (**E**) B, (**F**) myeloid, (**G**) NK, and (**H**) T cell proportions. In this and all subsequent figures unless otherwise indicated, boxplots depict the median via the thickest centre horizontal line, the first and third quartiles as the bottom and top of the box respectively, and 1.5x the interquartile range through whiskers; outliers are depicted as single points. Two-tailed t-test p-values are indicated for the self-reported ethnicity covariate in a model of log_10_(Proportion)∼Age+Sex+Individual_Self_reported_ethnicity. Gene expression UMAPs with cells coloured by log_2_(fold-enrichment) within a 500-cell neighbourhood, for (**I**) females versus males, (**J**) donors 50-77 years old versus other donors, and (**K**) SG_Malay donors relative to all other donors. Cell types of interest are indicated by dashed lines.

Based on the above cell types, we examined the relationship of age, female / male sex, and self-reported ethnicity with cell type proportions (cell counts relative to all PBMCs per donor, using a generalised linear model: log_10_(Proportion)∼Age+Sex+Self_reported_ethnicity). pDC proportions decreased with age (N=438, degrees of freedom (df)=430, t-value=-4.93, two-tailed t-test p-value=1.2e-06, **Figure 2B**). We confirmed that B cells were more abundant in females (N=562, df=554, t-value=-6.76, two-tailed t-test p-value=3.56e-11; **Figure 2C**), while NK cell proportions were higher in males (N=562, df=554, t-value=4.41, two-tailed t-test p-value=1.26e-05; **Figure 2D**). Thus, the AIDA dataset recapitulated known age^29^ and sex^30^ differences in circulating immune cells.

We then tested for differences across population groups in major cell type proportions, focusing on populations with ≥50 donors. One caveat for such analyses is that genetic variation and environmental factors are often confounded^31^. Here, the effects of self-reported ethnicity may represent a combination of genetic effects as well as correlated environmental and lifestyle factors (e.g., diet and geography), all of which can contribute to phenotypic variation. SG_Malay donors showed elevated B cell proportions relative to other population groups (log_10_(Proportion)∼Age+Sex+Individual_Self_reported_ethnicity (e.g., SG_Malay), N=562, df=558, t-value=3.02, two-tailed t-test p-value=0.00269; **Figure 2E**), while Thai donors had lower myeloid cell proportions (N=562, df=558, t-value=-3.21, two-tailed t-test p-value=0.00139; **Figure 2F**). NK cells were less abundant in SG_Indian donors (N=562, df=558, t-value=-4.57, two-tailed t-test p-value=5.91e-06; **Figure 2G**), while Korean donors had lower T cell proportions (N=562, df=558, t-value=-2.98, two-tailed t-test p-value=0.00298; **Figure 2H**). These results suggest systematic differences in major cell type proportions across population groups.

The proportions of peripheral blood cell types can serve as diagnostic markers for diseases, such as the ratio of monocytes to lymphocytes (for active tuberculosis^32^), relative monocyte proportion (for chronic myelomonocytic leukaemia and acute myeloid leukaemia^33^), as well as lymphocyte abundance (for lupus^14^). We found that the proportions of monocytes were lower on average in Thai donors than in donors of other population groups in both our scRNA-seq data (two-tailed Wilcoxon rank-sum p-value=3.08e-04, **Figure S1D**) as well as our complete blood count data (two-tailed Wilcoxon rank-sum p-value=3.28e-07, **Figure S1E**). Our findings suggest the importance of factoring in self-reported ethnicity in determining diagnostic baselines.

A major advantage of scRNA-seq is that differentially abundant cell populations can be characterised at high resolution. Through enrichment analysis of transcriptomic neighbourhoods, we identified more fine-grained trends than were apparent in the above analyses of major cell types. For example, although B cells were more abundant in female donors (**Figure 2C**), this trend was not uniform, and was more pronounced for naïve B cell populations (**Figure 2I**). In contrast, CD8^+^ T naïve cell populations were uniformly depleted in individuals ≥50-years-old (**Figure 2J**). The enrichment of gamma-delta T (γδT) cell populations in SG_Malay donors was also not uniform (**Figure 2K**). In these initial analyses, we examined the effects of sex, age, or self-reported ethnicity individually rather than in combination. Nevertheless, our results suggest that cell neighbourhood analyses illuminate biological differences not apparent at coarser resolutions.

### Cell type annotation and transcriptomic gradients

To interrogate more granular cell identities, we analysed three broad cell populations – B (**Figure 3A**), pDC and myeloid (**Figure 3B**), and ILC, NK, and T (**Figure 3C,D**) – separately. For each population, we performed feature selection, data integration, and sub-clustering independently to utilise features relevant for distinguishing the cell subtypes of interest^34^ (<u>Methods</u>, **Figure 3A-D,S2A,B**). We performed one round of sub-clustering for the first two populations. For ILC, NK, and T cells, after one round of sub-clustering, we separated CD4^+^ T and double-negative T (dnT) cells (**Figure 3C**) from all other ILC, NK, and T cells (**Figure 3D**), and integrated and re-clustered these two groups separately. All sub-clusters of all cell populations were then annotated based on marker genes (**Figures S2A,B**, **<u>Table S2</u>**) and presence of TCR barcodes (**Figure S1B**). We defined cluster identities as descriptions of individual sub-clusters, and cell subtypes as the manual merging of these sub-clusters into known subtypes.

**Figure 3:**
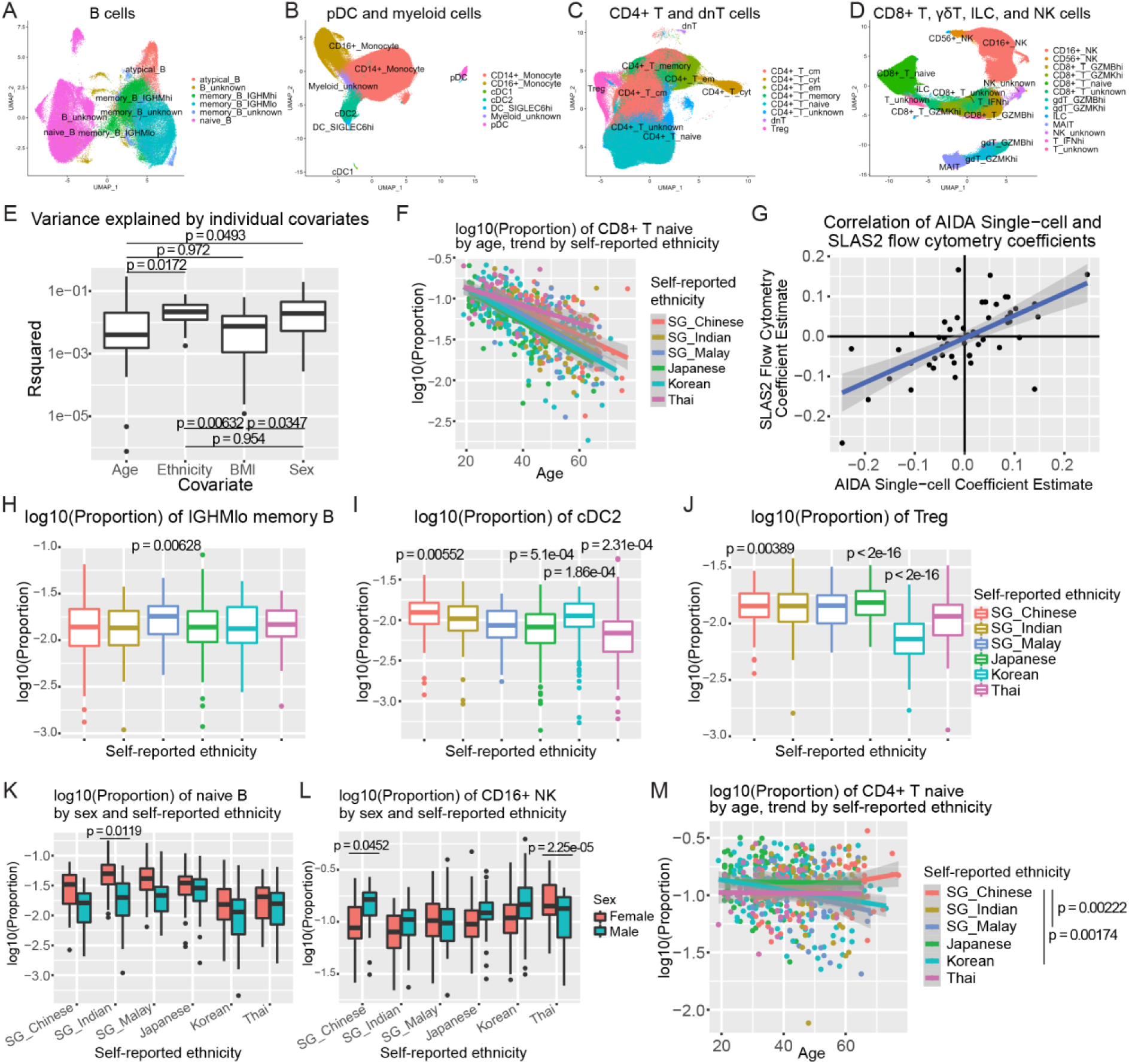
Relationships of human diversity with cell subtype proportions. Gene expression UMAPs depicting (**A**) B, (**B**) pDC and myeloid, (**C**) CD4^+^ T and dnT, and (**D**) CD8^+^ T, γδT, ILC, and NK sub-clusters labelled by cell subtype. (**E**) Boxplots depicting the variance in cell subtype proportions in Singapore donors explained by age, self-reported ethnicity, BMI, or sex when examined individually, annotated with all pairwise two-tailed Wilcoxon rank-sum p-values. Scatterplots depicting (**F**) CD8^+^ T naïve cell proportions against donor age for all AIDA donors, and (**G**) linear regression coefficient estimates for self-reported ethnicity in the SLAS-2 dataset (y-axis) versus the AIDA dataset (x-axis). Each point in (**G**) represents a combination of one of 18 cell subtype proportions regressed against one of the 3 Singapore self-reported ethnicities (log_10_(Proportion)∼Age+Sex+ Individual_Self_reported_ethnicity). Boxplots depicting cell subtype proportions for (**H**) *IGHM*^lo^ memory B, (**I**) cDC2, and (**J**) regulatory T (Treg) cells across all population groups. Boxplots depicting (**K**) naïve B and (**L**) CD16^+^ NK proportions across self-reported ethnicity and female / male sex. (**M**) Scatterplot depicting CD4^+^ T naïve proportions against donor age. Scatterplots are overlaid with linear regression lines for their respective data points; grey bands indicate the 95% confidence intervals. Two-tailed t-test p-values in (**H-J**) pertain to the self-reported ethnicity covariate in a model of log_10_(Proportion)∼Age+Sex+Individual_Self_reported_ethnicity. Two-tailed t-test p-values adjacent to lines indicate comparisons of two population groups in (**M**), and pertain to the interaction terms between sex and individual population groups in (**K,L**).

We identified rare cell subtypes such as dnT (0.04% of all cells), cDC1 (0.04%), and atypical B^35^ (0.4%) cells (**Figure 3A-C,S2A-C**). We further identified rare cluster identities, such as *SCART1*^hi^ ILC (0.02%) and *XCL1*^hi^ ILC (0.03%) (**Figure S2D**). The identification of these rare cell populations attests to the resolution of our scRNA-seq atlas.

To complement this discrete, categorical approach towards cell annotation, we examined continuous transcriptomic gradients. These included an *IGHM* gradient in memory B cells^36^ (**Figure S2E**); opposing *GZMB* and *GZMK* gradients in both CD8^+^ T memory^37^ and γδT cells, with heighted *GZMB* levels marking more cytotoxic T cell subsets; as well as opposing *FCER1G* and *KLRC2* gradients in CD16^+^ NK cells^38^ (**Figure S2F**). These continuous gene expression gradients may correlate with the effects of age, sex, and genetic variation in our cohort, which we describe below.

### Relationships of human diversity with cell subtype proportions

Haematological properties and immune cell subtype proportions are of broad interest as disease markers^14,39^, but these may be confounded by patient demographics. We therefore investigated the relationships of human diversity with cell subtype proportions. As a control, we confirmed that monocyte proportions in our scRNA-seq dataset were consistent with matched complete blood counts (**Figures S3A-C**), indicating that cell type proportion inferences from our scRNA-seq data were rooted in actual haematological proportions.

We first evaluated the relative impact of human demographics, by analysing the correlation of cell subtype proportions with covariates: age, BMI, self-reported ethnicity, or sex. For this analysis, we focused on donors profiled in Singapore to minimise the influence of technical variation across study sites. We examined the variance explained by a covariate of interest (<u>Methods</u>; **Figure 3E**). The highest variance explained for any human diversity-cell subtype combination was the decrease of CD8^+^ T naïve cell proportions with age (R-squared=0.290, N=200, df=198, t-value=-9.00, two-tailed t-test p-value<2e-16), consistent with previous reports (**Figure 3F**)^40,41^. More broadly, the proportions of multiple cell subtypes were significantly correlated with age. For example, CD4^+^ T cytotoxic cell proportions increased with age (N=501, df=499, t-value=4.86, p-value=1.57e-06 for a model of log_10_(Proportion)∼Age for all AIDA donors; **Figure S3D**); cytotoxic CD4^+^ T cells have been of interest for their heightened abundance in supercentenarians^42^. Overall, however, self-reported ethnicity and sex each explained more variance in subtype proportions than age or BMI (N=22, pairwise two-tailed Wilcoxon rank-sum p-values<0.05; **Figure 3E**).

To corroborate the associations between self-reported ethnicity and immune subtype proportions, we analysed published flow cytometry data (**<u>Table S3</u>**) from an independent cohort (Singapore Longitudinal Aging Study wave-2 (SLAS-2)^43^, <u>Methods</u>). First, we confirmed that MAIT cells were significantly higher in proportions in SG_Chinese than SG_Indian donors in both datasets (AIDA: SG_Indian coefficient estimate (versus SG_Chinese)=-0.315, N=198, df=193, t=-4.71, two-tailed t-test p-value=4.67e-06; SLAS-2: SG_Indian coefficient estimate (versus SG_Chinese)=-0.274, N=814, df=809, t=-3.46, two-tailed t-test p-value=5.7e-04; **Figure S3E**). Next, we examined effect size concordance of 54 coefficient estimates of self-reported ethnicity from linear models for cell subtype proportions (log_10_(Proportion)∼Age+Sex+Individual_Self_reported_ethnicity, i.e., SG_Chinese, SG_Malay, or SG_Indian). Effect sizes were well-correlated between AIDA and SLAS-2 (Pearson correlation r=0.652, N=54, df=52, t=6.20, two-tailed t-test p-value=9.41e-08, **Figure 3G**). This concordance across two modalities and independent cohorts supports our findings of self-reported ethnicity-associated cell subtype signatures in circulating immune cells.

We found numerous examples of self-reported ethnicity being associated with differential cell subtype proportions when we examined all AIDA population groups with ≥50 donors. Relative to all PBMCs per donor, *IGHM*^lo^ memory B proportions were higher in SG_Malay donors than in other population groups (log_10_(Proportion)∼Age+Sex+Individual_Self_reported_ethnicity (e.g., SG_Malay), N=562, df=558, t-value=2.74, two-tailed t-test p-value=0.00628; **Figure 3H**), while cDC2 proportions were lower in Thai donors (N=560, df=556, t-value=-3.71, two-tailed t-test p-value=2.31e-04; **Figure 3I**). Most strikingly, we found much lower regulatory T (Treg) proportions in Korean donors (N=562, df=558, t-value=-14.8, two-tailed t-test p-value<2e-16; **Figure 3J**). This effect remained when we controlled for possible differences in T cell proportions across population groups by assessing Treg proportions relative to CD4^+^ T cells (t-value=-14.4, two-tailed t-test p-value<2e-16, **Figure S3F**). We also observed a similar result when we re-clustered and re-annotated cells following data integration using a different algorithm (Harmony^44^; t-value=-14.4, two-tailed t-test p-value<2e-16, **Figure S3G**). We provide the associations of cell subtype proportions with self-reported ethnicity, controlling for age and sex, as a human diversity reference for the healthy ranges of immune cell subtypes across diverse population groups (**<u>Table S4</u>**), which may be incorporated as healthy baselines for diagnostics.

We hypothesised that interactions between self-reported ethnicity, age, and sex could affect cell subtype proportions. We extended the above linear models to include all pairwise interaction terms and found several examples. Interactions between sex and self-reported ethnicity could exacerbate female-male cell proportion differences: although naïve B cells were typically less abundant in males versus females, this disparity was greatest in SG_Indian donors (N=559, df=552, t-value=-2.52, two-tailed t-test p-value=0.0119 for interaction term in a model of SG_Indian donors versus all other donors; **Figure 3K**). In contrast, while CD16^+^ NK proportions were typically higher in males, this effect was not present in Thai donors (N=562, df=555, t-value=-4.27, two-tailed t-test p-value=2.25e-05 for interaction term in a model of Thai donors versus all other donors; **Figure 3L,S4A**). Similar to our Treg analyses above, we recapitulated these interactions after cell type re-annotation following data integration using the Harmony^44^ algorithm (naïve B: t-value=-2.51, two-tailed t-test p-value=0.0125, **Figure S4B**; CD16^+^ NK: t-value=-4.27, two-tailed t-test p-value=2.27e-05, **Figure S4C**). These results raise the intriguing possibility of cell type-specific differential sex hormone activity or sex chromosome gene regulation across population groups.

We also identified interactions between self-reported ethnicity and age: Korean and SG_Malay donors showed sharper decreases in CD4^+^ T naïve cell proportions with increasing age compared to SG_Chinese donors (N=562, df=543; t-value=-3.15, two-tailed t-test p-value=0.00174 for the age-Korean interaction; t-value=-3.07, two-tailed t-test p-value=0.00222 for the age-SG_Malay interaction, both compared against SG_Chinese donors; **Figure 3M,S4A**). Reduced CD4^+^ T naïve cell levels have been reported in hepatitis C virus-infected patients^45^ and SLE patients^14^. Given this disparity in CD4^+^ T naïve proportions across population groups and ages, these dimensions of human diversity need to be considered when assessing reference ranges and diseases where perturbed CD4^+^ T naïve cell proportions can serve as a biomarker.

### Single-cell signatures of human diversity

We harnessed the resolution afforded by our scRNA-seq atlas by investigating for single-cell signatures of human diversity in multiple cell populations (B; pDC and myeloid; CD4^+^ T and dnT; CD8^+^ T, γδT, ILC, and NK). We used MiloR^46^, implementing a model that incorporated multiple covariates (self-reported ethnicity, age, and female / male sex) for differential abundance testing of MiloR cell neighbourhoods in gene expression space. As a control, we examined cell neighbourhood abundances in males versus females. In males, the majority of naïve B cell neighbourhoods were depleted (MiloR neighbourhoods with MiloR spatial FDR<0.1, **Figure S5A**), while numerous CD16^+^ NK cell neighbourhoods were enriched (spatial FDR<0.1, **Figure 4A,S5B**). This was consistent with our cell subtype analyses (**Figure 3K,L,S4B,C**) and the patterns of B and NK cell type abundance reported previously^30,47^. As an additional positive control, we examined the most male-enriched B cell neighbourhood (log_2_(fold-change)=2.45, spatial FDR=4.80e-75), and found upregulation of non-pseudoautosomal region Y-chromosome genes, including *RPS4Y1*, *EIF1AY*, and *DDX3Y* (**Figure S5C**).

**Figure 4:**
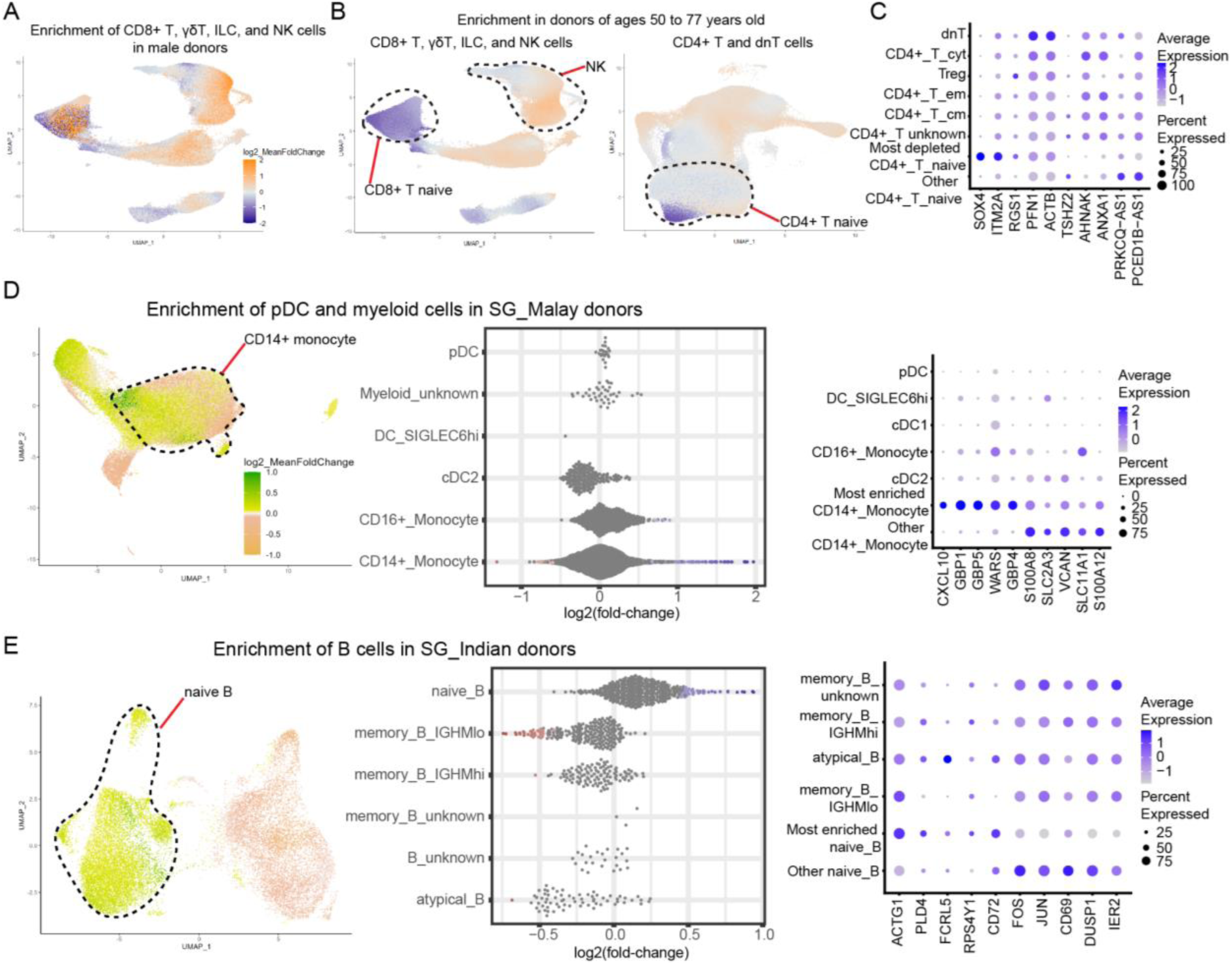
Single-cell signatures of human diversity. Gene expression UMAPs depicting enrichment of CD8^+^ T, γδT, ILC, and NK cell neighbourhoods in (**A**) males versus females, and (**B**) (left) donors ≥50-years-old versus younger donors, as well as (**B**) (right) enrichment of CD4^+^ T and dnT cell neighbourhoods in donors ≥50-years-old versus younger donors. (**C**) Dot plot of top 5 upregulated and top 5 downregulated genes (as compared to all other CD4^+^ T naïve cells) of the most depleted CD4^+^ T cell neighbourhood in donors ≥50-years-old versus younger donors. (**D,E**) (Left) UMAPs depicting enrichment of cell neighbourhoods in a cell population, (middle) beeswarm plots depicting enrichment of cell neighbourhoods, and (right) dot plots of top 5 upregulated and top 5 downregulated genes (as compared to all other cells of the cell type of interest) of the most enriched cell neighbourhood in the cell population. (**D**) pDC and myeloid cell neighbourhood enrichment for SG_Malay donors; a CD14^+^ monocyte neighbourhood shows the highest enrichment. (**E**) B cell neighbourhood enrichment for SG_Indian donors; a naïve B neighbourhood shows the highest enrichment. For UMAPs depicting cell neighbourhood enrichment, each cell is coloured by its log_2_(mean fold-change) value for all overlapping MiloR cell neighbourhoods that the cell was grouped in. Cell types of interest are indicated by dashed lines. In (**A-C**), the analysis was performed using all AIDA donors; orange hues indicate cell neighbourhood enrichment, while blue hues indicate cell neighbourhood depletion; darker hues correspond to higher magnitudes of enrichment or depletion, capped at log_2_(mean fold-change)=|2|. In (**D,E**), the analysis was performed on Singapore donors only; yellow-green hues indicate cell neighbourhood enrichment, while vermillion hues indicate cell neighbourhood depletion; darker hues correspond to higher magnitudes of enrichment or depletion, capped at log_2_(mean fold-change)=|1|. For beeswarm plots, each point corresponds to one cell neighbourhood; cell neighbourhoods are classified by the majority cell type annotation within the neighbourhood. Points coloured in red (depletion of neighbourhood for the dimension of human diversity of interest) and in blue (enrichment of neighbourhood) correspond to spatial FDR values<0.1.

We identified biases in cell neighbourhood abundance that were not evident in analyses of cell types or subtypes. For example, the lower abundance of B cells in males was not uniform across cell neighbourhoods, particularly for memory B cell neighbourhoods (**Figure S5A**). Similarly, NK cell neighbourhoods showed variable enrichment. While *STMN1*^hi^ NK cell neighbourhoods were enriched in males, CD56^+^ NK and IFN^hi^ CD16^+^ NK cell neighbourhoods were depleted in males (spatial FDR<0.1, **Figure S5B**). Furthermore, multiple *SOX4*^hi^ CD4^+^ T naïve cell neighbourhoods were enriched in females (spatial FDR<0.1, **Figure S5D**).

As additional controls, we examined the differential abundance of cell neighbourhoods in donors ≥50-years-old (∼25% of our cohort, **Figure 1D**) versus younger donors. We identified cell neighbourhoods that were enriched amongst CD16^+^ monocytes and depleted amongst pDCs and CD8^+^ T naïve cells in these older donors (spatial FDR<0.1, **Figure 4B,S6A,B**), concordant with our cell type and subtype analyses (**Figure 2B,3F**) and previous reports^48,49^.

We then identified age-associated single-cell signatures beyond those described in the literature. Multiple CD16^+^ NK cell neighbourhoods were significantly enriched in older donors (spatial FDR<0.1, **Figure 4B,S6B**), particularly cell neighbourhoods that were *FCER1G*^lo^ and *KLRC2*^hi^. Our analyses suggest an age bias in a subset of NK cells that were recently characterised at an scRNA-seq level as being adaptive NK cells^38^.

Furthermore, we found that a CD4^+^ T naïve cell neighbourhood with heightened *SOX4* expression (relative to other CD4^+^ T naïve cells) was the most depleted CD4^+^ T cell neighbourhood in donors ≥50-years-old (log_2_(fold-change)=-1.50, spatial FDR=1.61e-12, **Figure 4B,C,S6C**). This cell neighbourhood-based result was a more refined CD4^+^ T naïve aging signature than both that reported in the literature^47,49^ and seen in our cell subtype analyses (**Figure 3M**). This was also concordant with a finding from the OneK1K scRNA-seq study of European-ancestry donors in Australia, that cell counts of a particular CD4^+^ T cell subtype, with heightened *SOX4* expression and transcriptionally distinct from CD4^+^ T naïve and central memory cells, declined with age^17^. We note two coincidences in sex and age biases. CD4^+^ T naïve SOX4*^hi^* cell neighbourhoods were depleted in donors ≥50-years-old and enriched in females (spatial FDR<0.1, **Figure 4B,S5D,S6C**). In addition to the enrichment of *FCER1G*^lo^ *KLRC2*^hi^ CD16^+^ NK cell neighbourhoods in donors ≥50-years-old (spatial FDR<0.1, **Figure S6B**), multiple NK cell neighbourhoods were enriched in males (spatial FDR<0.1, **Figure S5B**). These results suggest the importance of considering multiple dimensions of human diversity in cellular and molecular analyses.

We tested for hitherto unexplored single-cell signatures of self-reported ethnicity, focusing on Singapore donors to minimise confounding by technical variation across study sites. SG_Malay donors showed enrichment of a CD14^+^ monocyte cell neighbourhood (log_2_(fold-change)=1.97, spatial FDR=2.28e-07; **Figure 4D**) with heightened expression of *GBP1, GBP4, GBP5,* and *WARS* (interferon-induced genes^50,51^) and *CXCL10* (an interferon-induced chemokine) relative to other CD14^+^ monocytes (**Figure 4D**). The majority of γδT *GZMB*^hi^ cell neighbourhoods (**Figure 2K,S2C**) were enriched in SG_Malay donors (spatial FDR<0.1, **Figure S6D**). SG_Indian donors showed enrichment of a naïve B cell neighbourhood (log_2_(fold-change)=0.938, spatial FDR=1.00e-03, **Figure 4E**) with heightened expression of *ACTG1 and PLD4*, and lowered *CD69* levels (**Figure 4E**). *PLD4* is a marker gene for IGM^hi^ transitional B cells^52^, while CD69 is an early marker of lymphocyte activation^53^, suggesting that cells in this SG_Indian-enriched neighbourhood may be in a progenitor-like state.

These examples highlight sex, age, and self-reported ethnicity differences discernible only at the level of cell neighbourhood abundance rather than at the resolution of cell types or subtypes. Collectively, through these analyses, we have identified single-cell signatures of human diversity, which point to fundamental differences in immune cell phenotypes across diverse donors. These results demonstrate the importance of considering sex, self-reported ethnicity, and age in the inference of disease signatures to avoid the confounding of disease with diversity.

### Molecular variation across population groups

Having elucidated numerous cell type, subtype, and neighbourhood signatures associated with multiple dimensions of human diversity, we investigated the influence of self-reported ethnicity on cell subtype-specific gene expression. We used edgeR^54^ to test for population group-specific differentially expressed genes (DEGs) based on pseudobulk transcriptomes aggregated per cell subtype per donor (<u>Methods</u>). We incorporated age, sex, and scRNA-seq experimental batch as covariates in this analysis, and focused on the Singapore donors to minimise the confounding of self-reported ethnicity differences with technical variation across study sites.

We identified genes that showed a consistent pattern of self-reported ethnicity-associated expression across most subtypes. For example, *UTS2* (a gene encoding a cyclic peptide with strong vasoconstrictor activity^55^) was differentially expressed in the highest number of cell subtypes (16 subtypes, alongside *KANSL1*, *NSF*, and *PPDPF*; **<u>Table S5</u>**) out of all genes tested. *UTS2* showed consistent upregulation in SG_Chinese donors (edgeR log2(fold-change) 1.01 to 1.60) and downregulation in SG_Indian donors (edgeR log2(fold-change) - 2.86 to −0.918) across major immune cell types (FDR<0.05 for SG_Indian in 16 cell subtypes and SG_Chinese in 9 cell subtypes; **Figure 5A**, **<u>Table S5</u>**). This self-reported ethnicity-specific expression pattern was also observed in a microarray-based study of RNA extracted from whole blood, which ranked *UTS2* amongst the top significantly differential probe sets across the Singapore population groups^56^.

**Figure 5:**
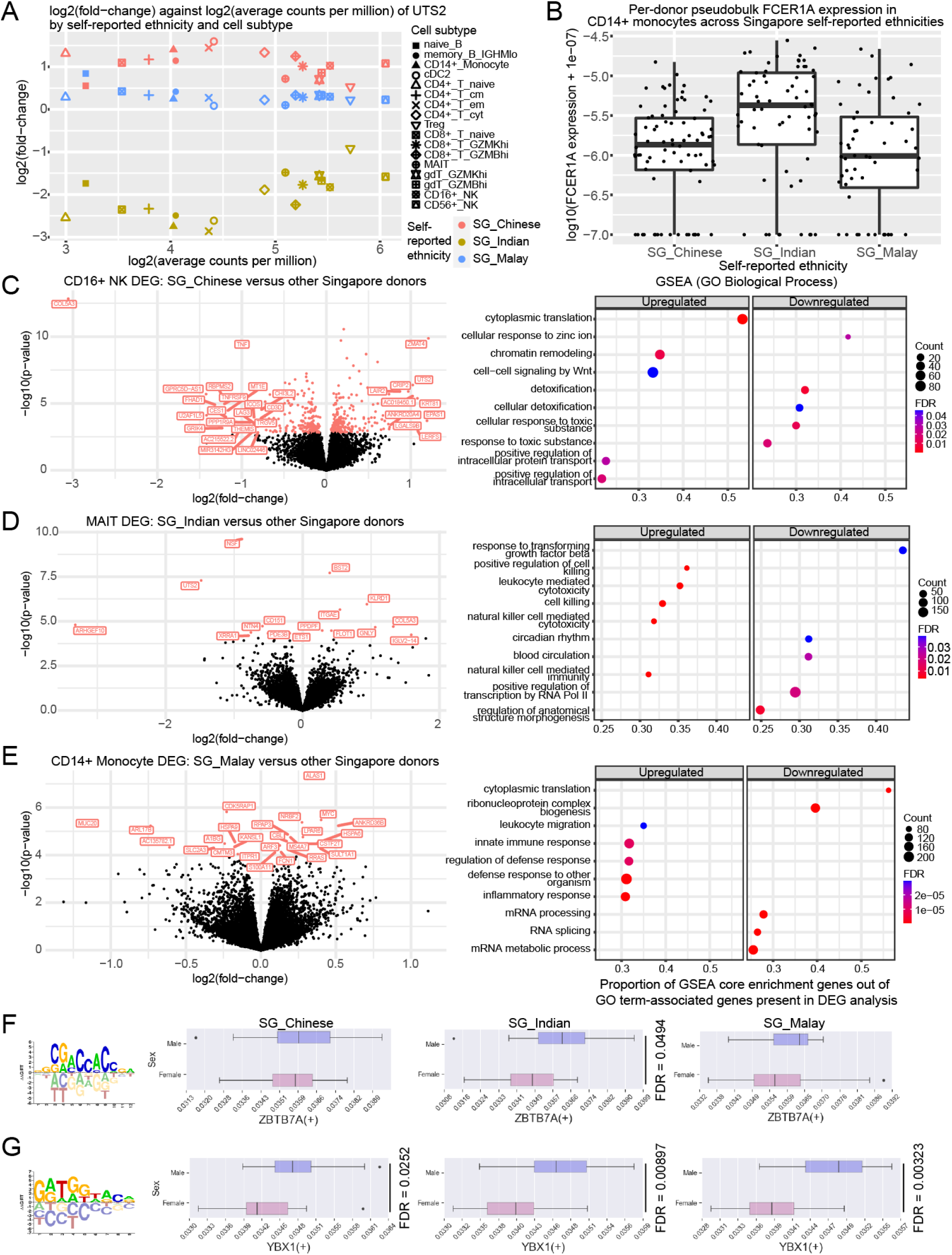
Self-reported ethnicity-associated variation in gene expression. (**A**) Scatterplot of edgeR-log_2_(fold-change versus other population groups) against edgeR-log_2_(average counts per million) of *UTS2* expression in cell subtypes for each Singapore self-reported ethnicity. (**B**) Boxplots of log_10_ transformation of per-donor pseudobulk *FCER1A* expression values normalised by total UMIs (with added pseudocount of 1e-07) in CD14^+^ monocytes across the Singapore self-reported ethnicities. Each dot represents a donor. Volcano plots of edgeR -log_10_(p-value) versus edgeR log_2_(fold-change) of differentially expressed genes in (**C**) (left) CD16^+^ NK cells in SG_Chinese donors, (**D**) (left) MAIT cells in SG_Indian donors, and (**E**) (left) CD14^+^ monocytes in SG_Malay donors, each versus all other Singapore donors. Points coloured in red have FDR<0.05, computed from Benjamini-Hochberg correction^57^ of p-values for genes analysed in the self-reported ethnicity-cell subtype combination. Points labelled with gene names have FDR<0.05 for (**C**-**E**), and also |log_2_(fold-change) values|≥0.75 for (**C**). (**C**-**E**) (Right) Gene set enrichment analysis (GSEA) dot plots of the top ∼5 (based on GSEA p-value) upregulated or downregulated (positive or negative enrichment score, respectively) Gene Ontology (GO) Biological Process gene sets. Dot size (“Count”) indicates number of core enrichment genes; dots are coloured by FDR (Benjamini-Hochberg-corrected^57^ p-values). (**F**,**G**) (Left to right) Transcription factor (TF) binding site motif from CIS-BP^58^, and boxplots depicting the distributions of the median TF regulon AUCell score across all cells of the subtype of interest per donor for each of the indicated Singapore population groups. (**F**) *ZBTB7A* (M02914_2.00) in regulatory T (Treg) cells; (**G**) *YBX1* (M04661_2.00) in CD4^+^ T effector memory (em) cells. FDR values of comparisons between males versus females in (**F**,**G**) were computed from Benjamini-Hochberg correction^57^ of all two-tailed Wilcoxon rank-sum p-values for a regulon across the SCENIC GRNBoost2 trial-AUCell analysis combinations. Boxplots depict the output from one SCENIC GRNBoost2 trial-AUCell analysis combination.

Furthermore, *FCER1A* was upregulated in SG_Indian donors (edgeR log_2_(fold-change)=1.45, FDR=2.92e-05) and downregulated in SG_Chinese (edgeR log_2_(fold-change)=-1.16, FDR=0.0108) in CD14^+^ monocytes (**Figure 5B**). *FCER1A* expression in monocytes has been implicated in risk of allergic disease, with a GWAS SNP for allergic disease risk, rs2427837, associated with *FCER1A* gene expression and protein levels in a Singaporean Chinese cohort^59^. Genetic variation associated with *FCER1A* may also be predictive for treatment response in adult East Asian patients with chronic hepatitis B^60^. The self-reported ethnicity-specific expression patterns of *FCER1A* may be explained in part by differences in the allele frequency of rs2427837 (chr1_159288755_G_A) across the Singapore population groups (**Figure S7A**). Differential expression of disease-associated genes across population groups, such as *FCER1A*, at a level akin to the magnitude of gene expression variation linked with eQTLs, may highlight genes of interest for investigating differential disease risk and susceptibility across population groups.

Across 21 cell subtypes and in comparison to the other Singapore self-reported ethnicities, we identified 1,915 DEGs for SG_Chinese donors and 1,968 for SG_Indian donors, but only 97 for SG_Malay donors (FDR<0.05, **<u>Table S5</u>**). The low number of SG_Malay DEGs may reflect the intermediate position of this population group in the genotype PCA between SG_Chinese and SG_Indian (**Figure 1E**), and suggests that the SG_Malay donor group may also occupy an intermediate position in gene expression space. In CD16^+^ NK cells, genes upregulated in SG_Chinese donors were enriched for a Wnt signalling-associated gene set (FDR<0.05, **Figure 5C**); Wnt signalling has been implicated in NK-cell differentiation and function^61^. For MAIT cells, SG_Indian donors showed upregulation of cytotoxicity-related gene sets and downregulation of a TGFβ-response gene set (FDR<0.05, **Figure 5D**), which is intriguing considering the lower proportions of these innate immune system-like^56^ cells in SG_Indian versus SG_Chinese donors (**Figure S3E**). SG_Malay donors showed upregulation of gene sets associated with inflammatory and host-pathogen defence responses in CD14^+^ monocytes (FDR<0.05, **Figure 5E**), which is concordant with the enrichment of CD14^+^ monocyte cell neighbourhoods with elevated expression of interferon-associated genes in SG_Malay donors (**Figure 4D**). These results suggest cell subtype-specific expression differences that may underlie variation in the activity levels of biological pathways across population groups.

We further hypothesised that differences in gene regulatory networks could contribute to expression variation across population groups. We used a SCENIC-based^62^ workflow to identify transcription factor regulons (comprising a transcription factor and its target genes) whose activities differed across dimensions of human diversity. For example, we observed that *ZBTB7A* regulon activity in Treg cells was heightened in male versus female donors of SG_Indian self-reported ethnicity (regulon-specific FDR<0.05, **Figure 5F**), but this sex difference was not apparent in other Singapore self-reported ethnicities. ZBTB7A may have both transcriptional activator and repressor properties^63^, and has been reported as a repressor of genes involved in glycolysis^64^ and foetal globin gene expression^65^. In contrast, *NFYC* did not show sex-specific regulon activity in Treg cells for any Singapore self-reported ethnicity (**Figure S7B**). We further identified a possible sex bias in *YBX1* regulon activity in CD4^+^ T effector memory (em) cells across all Singapore self-reported ethnicities (regulon-specific FDR<0.05, **Figure 5G**); YBX1 represses interferon gamma-induction of human major histocompatibility complex (MHC) class II genes^66^.

Together, these analyses suggest a possible relationship between self-reported ethnicity (including its correlated environmental and lifestyle factors) and cell type-specific molecular phenotypes, such as gene expression levels, as well as gene set and gene regulatory network activity. Such molecular variation may contribute to physiological and disease-related variation across population groups, such as variation in immune responses and haematological features.

### eQTL analyses identify population-specific functional variants and contextualise disease-associated loci

To investigate additional influences of human diversity on cell type-specific phenotypes, we performed pseudobulk eQTL analysis for 20 immune cell subtypes (<u>Methods</u>). Our cell subtype-specific eQTL analyses circumvent the confounding between differential cell proportions versus differential gene expression that eQTL analyses of bulk tissues (e.g., whole blood) suffer from^67^. We identified 11,431 unique genes with at least one cis-eQTL (within 1 Mb of the gene) with FDR<0.05 in a cell subtype (eGene), out of 12,187 unique autosomal genes analysed across all subtypes (**Figure 6A**). The number of eGenes discovered per cell subtype correlated with the number of donors analysed for the corresponding subtype (**Figure 6A,S8A**), with a median of 2,342 such eGenes per cell subtype (range: 366 to 6,444). We also implemented eigenMT^68^ for p-value correction from our eQTL analyses to nominate one lead SNP per gene (**<u>Table S6</u>**).

**Figure 6:**
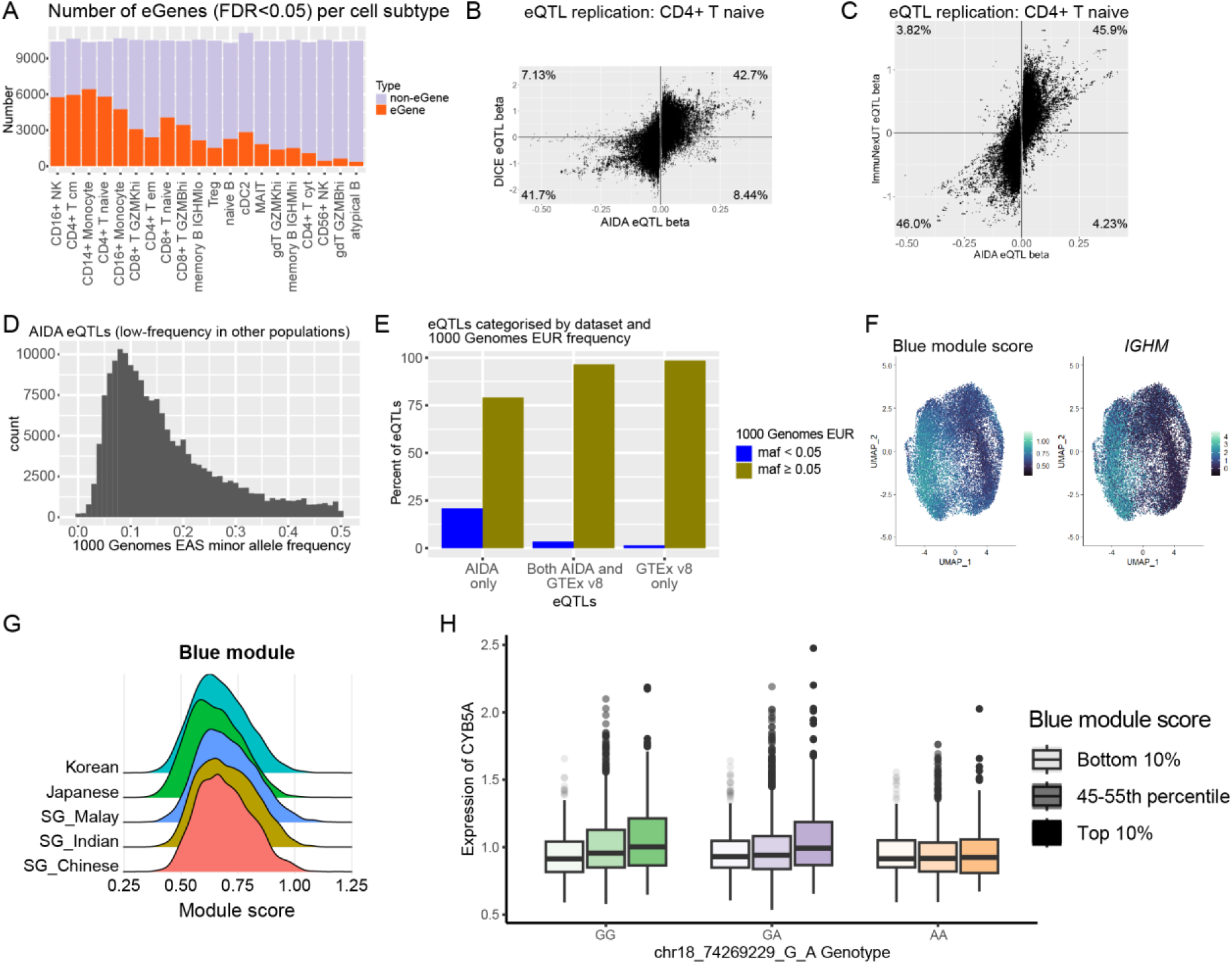
Population-specific and context-dependent eQTL effects. (**A**) Bar charts of numbers of eGenes (Matrix eQTL FDR<0.05 for each cell subtype) and non-eGenes per cell subtype, ordered by number of donors analysed per cell subtype. Scatterplots of (**B**) DICE^69^ (y-axis) and (**C**) ImmuNexUT^70^ (y-axis) versus AIDA (x-axis) CD4^+^ T naïve eQTL effect size (beta) values of SNP-gene pairs with AIDA eQTL FDR<0.05 per cell subtype. Percentages of all SNP-gene pairs that lie within a quadrant are indicated. (**D**) Histogram of minor allele frequencies (maf) in the 1000 Genomes East Asian (EAS) super-population, for AIDA eQTLs that were low frequency (maf 1%-5%) or rare (maf<1%) in at least one of the 1000 Genomes African (AFR), Admixed American (AMR), or European (EUR) super-populations. (**E**) Bar charts of (left to right) eQTLs identified in AIDA only, eQTLs identified in both AIDA and GTEx v8 whole blood^18^, and eQTLs identified in GTEx v8 whole blood only; SNPs examined were present in both datasets. Bars depict percentages of eQTLs in the respective category with maf≥0.05 or maf<0.05 in the 1000 Genomes EUR super-population. (**F**) Feature plots depicting (left) blue module scores and (right) *IGHM* expression coloured on gene expression UMAPs of AIDA *IGHM*^hi^ and *IGHM*^lo^ memory B cells. (**G**) Ridge plots depicting the distributions of blue module scores across AIDA population groups. (**H**) Boxplots depicting *CYB5A* expression in AIDA *IGHM*^hi^ and *IGHM*^lo^ memory B cells, categorised by the magnitude of a cell’s blue module score, and by donor genotype of the chr18_74269229_G_A locus.

We corroborated our eQTLs using datasets from the DICE project, which profiled 91 healthy donors in the San Diego area, California, USA^69^. AIDA eQTLs (FDR<0.05 per subtype) were 84.1% to 87.1% concordant in effect size direction for 5 immune cell subtypes (CD4^+^ T naïve, naïve B, CD14^+^ monocyte, CD8^+^ T naïve, and CD16^+^ NK; **Figure 6B,S8B**). This indicated a substantial degree of replication in eQTL identification for the AIDA eQTL set. We observed an enhancement in eQTL replication when comparing AIDA eQTLs (FDR<0.05 per subtype) against ImmuNexUT eQTLs (derived from purified immune cell subsets from 416 Japanese donors) for the same 5 cell subtypes^70^ (92.0% to 93.9% concordance, **Figure 6C,S8C**). This improvement in replication may be attributable to the larger size of the ImmuNexUT cohort versus the DICE cohort, as well as the closer genetic similarity between the AIDA and ImmuNexUT cohorts than that for the AIDA and DICE cohorts.

We examined the allele frequencies of AIDA eQTLs present in the 1000 Genomes Phase 3 (ENSEMBL release 105) dataset^71^. 6.94% of such variants were low frequency or rare (minor allele frequency (maf) 1%-5%, and maf<1%, respectively) in each of the African (AFR), Admixed American (AMR), and European (EUR) super-populations. 2.24% of AIDA eQTL variants were entirely absent in the EUR super-population. 31.6% of AIDA eQTL variants were low-frequency or rare in at least one of the aforementioned super-populations, with many of these being common in the East Asian (EAS) and South Asian (SAS) super-populations (**Figure 6D,S8D**). In addition, we compared the set of AIDA eQTLs with FDR<0.05 (Benjamini-Hochberg-adjusted^57^ p-values) across all tests in all cell subtypes against eQTLs (Benjamini-Hochberg-adjusted^57^ p-value<0.05) found in the GTEx v8 whole blood dataset^18^ (**Figure 6E**), focusing on SNPs tested in both studies. Of the variants identified as eQTLs only in AIDA, 20.9% were present at maf<0.05 in the 1000 Genomes EUR super-population, but were common in the AIDA cohort (**Figure 6E**). This was much higher than the corresponding proportion (3.46%) for eQTLs found in both AIDA and the GTEx v8 whole blood dataset (**Figure 6E**). These allele frequency differences are consistent with the existence of functional population-specific variants, and demonstrate the importance of studying diverse populations for characterising the full spectrum of genetic variants relevant to humanity.

We leveraged the single-cell resolution of our dataset to elucidate context-dependent eQTL effects (<u>Methods</u>), which can go beyond cell type-specific analyses to pinpoint cellular mechanisms and cell states modulating gene expression variation^72^. We modelled such cellular contexts using gene modules identified through gene-gene correlation analyses, such as a gene module (“blue”) corresponding to the *IGHM* gradient in memory B cells (**Figure 6F,G,S2E**), which may relate to B cell activation^73^. We identified 7,597 (13.9%) out of 54,798 SNP-gene pairs tested that showed eQTL effects dependent on the blue module cellular context (FDR<0.05 from Benjamini-Hochberg-adjusted^57^ p-values). For example, the impact of rs7239151 (chr18_74269229_G_A) on *CYB5A* expression varied with the magnitude of a cell’s blue module score. Higher module scores correlated with higher *CYB5A* expression for the GG genotype, but not for the AA genotype (**Figure 6H**), indicative of the modulation of variant effects by B cell activation status. 7.25% of variants showing context-dependent eQTL effects are common in AIDA but low-frequency or rare in the 1000 Genomes EUR super-population (**Figure S8E**), suggesting the presence of context-dependent effects that can only be discovered by studying diverse population groups.

In parallel, we performed colocalisation analyses for the AIDA eQTLs with immune-related disease GWAS featuring Asian cohorts (**<u>Table S7</u>**). We identified 1,025 cases of high posterior probabilities (PP) of colocalisation across 20 cell subtypes (colocalisation PP>0.8; <u>Methods</u>; **<u>Table S8</u>**). For example, we identified rs56750287, an AIDA eQTL for *ORMDL3* in CD8^+^ T *GZMK*^hi^ cells (N=458, df=431, t-value=-17.7, two-tailed t-test p-value=4.20e-53) and a trans-ancestry GWAS variant for rheumatoid arthritis^74^ (GWAS p-value=7.1e-13; colocalisation PP=0.810, **Figure 7A**). Variants affecting *ORMDL3* expression have been linked with inflammatory diseases such as childhood-onset asthma^75^.

**Figure 7:**
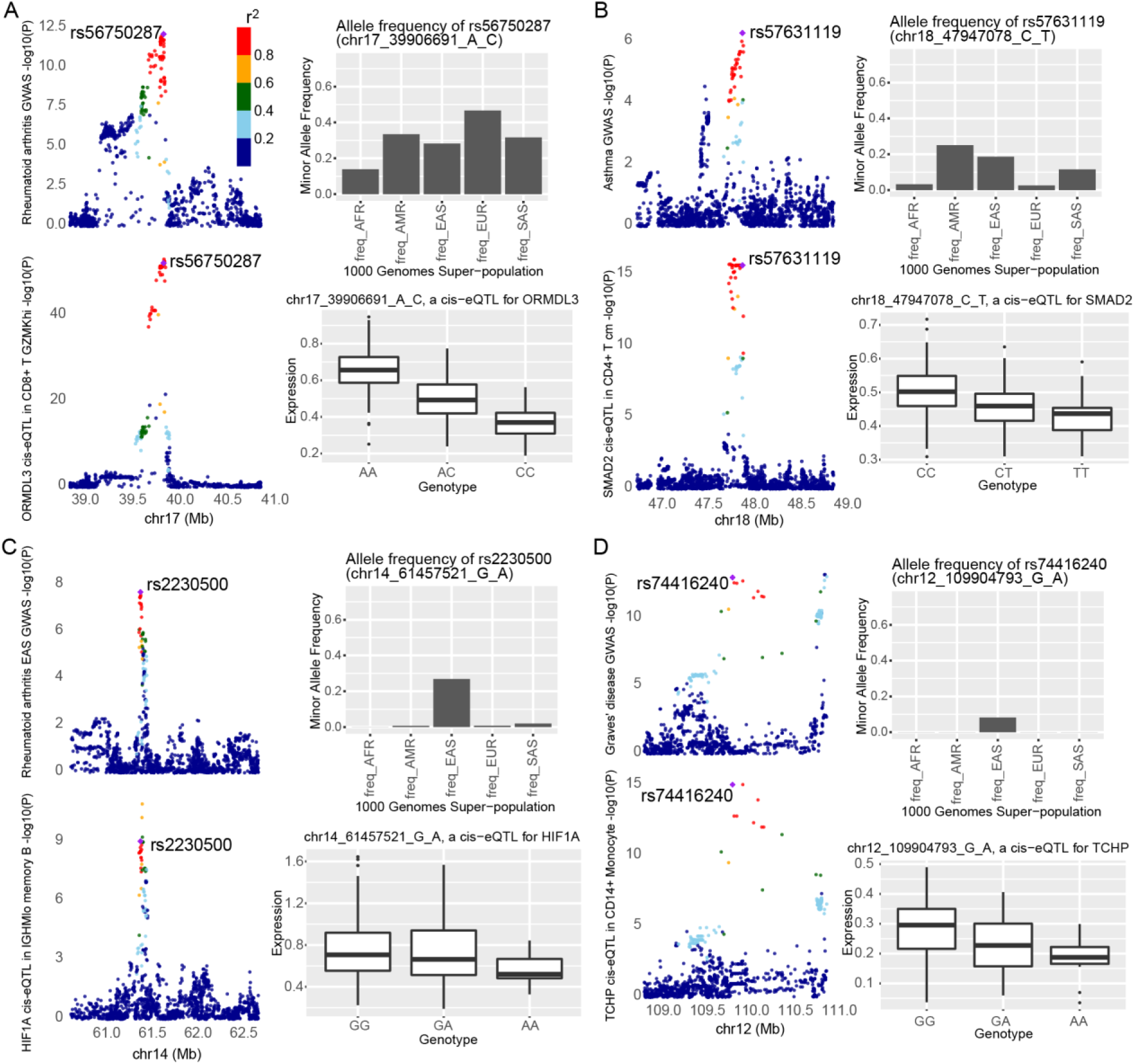
eQTL analyses contextualise disease-associated loci. Locus plots in (**A**-**D**) of variants investigated in GWAS (top left) and AIDA eQTL analyses (bottom left). (Top right) Minor allele frequencies of variant of interest in the 1000 Genomes super-populations (African (AFR), Admixed American (AMR), East Asian (EAS), European (EUR), and South Asian (SAS)). (Bottom right) Boxplots of eGene expression in the implicated cell subtype (y-axis) against AIDA donor genotypes (x-axis). (**A**) rs56750287 as an AIDA eQTL for *ORMDL3* in CD8^+^ T *GZMK*^hi^ cells and a rheumatoid arthritis trans-ancestry GWAS variant. (**B**) rs57631119 as an AIDA eQTL for *SMAD2* in CD4^+^ T cm cells and an asthma GWAS variant. (**C**) rs2230500 as an AIDA eQTL for *HIF1A* in *IGHM*^lo^ memory B cells and a rheumatoid arthritis EAS GWAS variant. (**D**) rs74416240 as an AIDA eQTL for *TCHP* in CD14^+^ monocytes and a Graves’ disease GWAS variant.

We then investigated possible population-specific causal variants, akin to the pathogenic transthyretin V122I missense variant found in patients of African ancestry^76^ and implicated in an under-diagnosed cause of heart failure amongst African American individuals. We identified numerous examples of population-specific variants in our colocalisation analyses across all major cell populations and spanning multiple diseases. rs57631119, an eQTL for *SMAD2* in CD4^+^ T cm cells (N=461, df=434, t-value=-8.54, two-tailed t-test p-value=2.19e-16) and a GWAS variant for asthma^77^ (GWAS p-value=5.29e-07; colocalisation PP=0.951), was a low-frequency (1-5%) variant in the 1000 Genomes AFR and EUR super-populations but common in all other super-populations (**Figure 7B**). TGFβ-SMAD2 signalling is active in the airways of asthmatic individuals^78^. rs2230500, an eQTL for *HIF1A* in *IGHM*^lo^ memory B cells (N=435, df=408, t-value=-6.28, two-tailed t-test p-value=8.77e-10) and also a variant identified in both a trans-ancestry and in particular an East Asian ancestry GWAS for rheumatoid arthritis^74^ (GWAS p-value=2.03e-08; colocalisation PP=0.864), was rare (<1% minor allele frequency) in non-EAS and non-SAS super-populations (**Figure 7C**). HIF1A has been implicated in angiogenesis and inflammatory activity in the synovium tissue of patients with rheumatoid arthritis^79^.

Furthermore, a variant identified only in the EAS super-population, rs74416240, was an AIDA eQTL for *TCHP* in CD14^+^ monocytes (N=460, df=433, t-value=-8.36, two-tailed t-test p-value=8.48e-16) as well as a variant implicated in Graves’ disease^80^ (GWAS p-value=8.61e-14; colocalisation PP=0.983, **Figure 7D**). Though little is known about *TCHP* in disease, with no disease reports in OMIM^81^ or UniProt^82^, this result suggests that *TCHP* may be a candidate for functional investigation for Graves’ disease.

Collectively, our genetic analyses provide a rich resource for understanding cellular and molecular mechanisms that may underlie disease risk, disease biology, and human phenotypic variation. Our QTLs facilitate the prioritisation of loci, genes, cell subtypes, and cellular contexts, including those relating to population-specific variants, for variant analysis and interpretation.

## Discussion

We have generated and assembled an scRNA-seq immune cell atlas from healthy donors spanning diverse population groups across 5 countries, which is a resource that is one of the largest healthy blood datasets in terms of number of cells, and also the most diverse in terms of number of population groups. We observed a substantial influence of human diversity on cellular and molecular traits. In our data, the impact of self-reported ethnicity on cell subtype proportions was comparable to that of female / male sex, and explained more variance than age or BMI. We found that the effects of age and sex could be modulated by self-reported ethnicity, highlighting the importance of studying these factors in combination. We identified scRNA-seq signatures of human diversity, including enrichment or depletion discernible only at the resolution of cell neighbourhoods, rather than cell types or subtypes. Human diversity-associated variation in cell subtype proportions, cell neighbourhood abundance, and cell type-specific gene expression overlapped with traits relevant to disease signatures and diagnostics.

These included population-specific functional variants, including context-dependent eQTLs, which could only be elucidated and characterised by studying diverse cohorts. Our findings underline how studying human diversity is both a pressing equity issue and of scientific importance. All the dimensions of human diversity we investigated influence fundamental phenotypes, and would ideally be incorporated in scRNA-seq reference atlases to facilitate accurate inferences from comparisons involving disease datasets.

Since genetic variation and environmental factors are often confounded^31^, the self-reported ethnicity associations we discovered are not necessarily genetic effects. Such associations may represent the combined effects of systematic genetic and environmental differences between populations, including variation in diet and geography. Further research is needed in cohorts with comprehensive metadata on environment and lifestyle, to begin to understand the individual impact of these factors. Regardless of their aetiology, these self-reported ethnicity-associated cellular and molecular profiles can contribute towards defining healthy baselines important for disease diagnoses, and facilitate the analysis of human physiology and pathophysiology.

Technical variation across experimental batches and study sites can introduce statistical biases confounded with the biological variation of interest. We adopted a suite of experimental and computational techniques to ameliorate against this, by harmonising experimental workflows and unifying data analysis across our study sites. In addition, we confirmed that key results remained consistent across two data integration methods (**Figure 3J-L,S3G,S4B,C**). To further reduce the impact of site-specific technical biases on our analyses of self-reported ethnicity, we performed internal comparisons within population groups, such as female-male or age differences within each population group, and focused several analyses on our Singapore cohort, for whom SG_Chinese, SG_Malay, and SG_Indian donors were batch-randomised. Lastly, we validated our findings in independent cohorts and datasets beyond our AIDA scRNA-seq atlas. These strategies may be relevant for future multi-national single-cell studies of diverse cohorts.

In our scRNA-seq atlas, *UTS2*, *KANSL1*, and *ANKRD36B* were the most commonly upregulated genes across multiple cell subtypes for SG_Chinese, SG_Indian, and SG_Malay donors, respectively (**<u>Table S5</u>**). *UTS2* was associated with multiple traits in a transcriptome-wide association study of the UK Biobank cohort^83^, including basal metabolic rate, body fat percentage, and diastolic blood pressure. Notably, the Singapore 2022 National Population Health Survey^84^ reported differences across Singapore self-reported ethnicities in the crude prevalence of abdominal obesity, hyperlipidaemia, type 2 diabetes, and hypertension, motivating future studies of the relationships between single-cell molecular variation and phenotypic variation across population groups.

Our findings of differential cell subtype proportions, such as the lower Treg proportions in Korean donors (**Figure 3J,S3F,G**), may have implications for our understanding of population-specific disease susceptibility. Treg cell depletion has been observed in the pathogenesis of autoimmune diseases^85^, motivating further studies of autoimmune disease prevalence in Korean populations, and more broadly comparative studies of disease prevalence across ancestries. For example, a study focusing on Manhattan, New York, USA, found that the prevalence of lupus was higher among non-Hispanic Asian women than non-Hispanic white women^86^. The combined analysis of detailed immune phenotypes elucidated from scRNA-seq datasets with studies of disease prevalence across diverse population groups can help nominate mechanisms of interest for understanding and treating diseases.

Given the resolution of our dataset, we could identify cell neighbourhood differences that were not apparent at the coarser cell type or subtype levels, such as the enrichment of CD14^+^ monocyte neighbourhoods with heightened expression of interferon-induced genes in SG_Malay donors (**Figure 4D**). Our findings both encourage further studies on the possibility of differential cell activities across population groups as well as warn against interpreting cell neighbourhood enrichment without accounting for human diversity. Such population-specific features should be factored into the analysis of disease scRNA-seq datasets. Comparisons of disease datasets against scRNA-seq reference atlases can nominate disease-associated cell neighbourhoods. However, if the reference atlases are not well-matched for dimensions of human diversity, such differentially abundant cell populations may be confounded with single-cell signatures specific to donor or patient demographics. With the possible applications of scRNA-seq technologies in precision medicine^39^, scRNA-seq reference atlases should be diverse from their inception to maximise the global benefit to all population groups.

## Acknowledgements

We would like to thank all donors and participants in the studies constituting the Asian Immune Diversity Atlas. This project was funded by grant number CZF2019-002446 (S.P., W-Y.P., J.W.S., and J.C.C.) from the Chan Zuckerberg Foundation, and grant numbers 2020-224570 (S.P., V.C., P.M., and P.P.M.), 2021-240178 (S.P., W-Y.P., J.W.S., J.C.C., V.C., P.M., and P.P.M.), and 2023-330381 (K.H.K.) from the Chan Zuckerberg Initiative DAF, an advised fund of Silicon Valley Community Foundation. The Singapore donor samples were obtained through the Health for Life in Singapore (HELIOS) Study (Lee Kong Chian School of Medicine, Nanyang Technological University (NTU); National Healthcare Group, Singapore; Imperial College London). We would like to express our gratitude to participants of the HELIOS study and the HELIOS operation team for recruitment, organisation, and data and sample collection, including Yoke Yin Terry Tong, Swat Kim Kerk, Guo Liang Low, and Halimah Binte Ibrahim from the HELIOS Biobanking team. This study (NTU IRB: 2016-11-030) is supported by the Singapore Ministry of Health’s (MOH) National Medical Research Council (NMRC) under its OF-LCG funding scheme (MOH-000271-00) and intramural funding from Nanyang Technological University, Lee Kong Chian School of Medicine, and the National Healthcare Group. The SG10K_Health project is funded by the A*STAR Industry Alignment Fund (Pre-Positioning) (IAF-PP: H17/01/a0/007). This project was also supported by funding from the A*STAR IAF-PP: H18/01/a0/020 (S.P.), the Japan Ministry of Education, Culture, Sports, Science and Technology (MEXT) to RIKEN Center for Integrative Medical Sciences (IMS), the Thailand Program Management Unit for National Competitiveness Enhancement (PMU-C) (C10F650132) (V.C., P.M., M.P., and B.S.), and Mahidol University’s Basic Research Fund: fiscal year 2021 (BRF1-017/2564) (V.C. and B.S.). We would like to thank the A*STAR Singapore Immunology Network (SIgN) Flow Cytometry platform for enabling the flow cytometry data analysis in this publication. The flow cytometry data analysis was supported by the A*STAR Joint Council Office (grant number 1434M00115), and the flow cytometry platform was supported by the SIgN Immunomonitoring Platform (BMRC IAF grant number 311006 and BMRC transition funds number H16/99/b0/011) as well as the National Research Foundation (NRF) Immunomonitoring Service Platform grant (NRF2017_SISFP09). B.L. is supported by the Singapore Ministry of Education, under its Academic Research Fund Tier 1 (FY2023; 23-0434-A0001; 22-5800-A0001) and Tier 2 (MOE-T2EP30123-0015), the Precision Medicine Translational Research Programme Core Funding (NUHSRO/2020/080/MSC/04/PM), National University of Singapore (NUS) ODPRT Seed Funding, and NUS YLLSoM Seed Funding. Y.Tong is supported by the MGI IRP scholarship. We would like to thank Jennifer Zamanian, Jennifer Chien, and Jason Hilton from the Human Cell Atlas Lattice team (Stanford University) for their help with and work on data deposits and coordination for community access to scRNA-seq datasets. We would like to thank members of our laboratories, the Chan Zuckerberg Initiative Single-Cell Biology team, and Mohamad Amin Honardoost for feedback and helpful discussions. This publication is part of the Human Cell Atlas – www.humancellatlas.org/publications/.

## Author contributions

K.H.K. pre-processed and analysed data, contributed towards study design, and supervised single-cell data analysis. L.M.T., K.Y.H., Y.A., D.J, A.C., J.C., S.G., M.K., T.K., J.L., S.N., S.Sarkar, N.T., and P.N.V. performed scRNA-seq experiments. K.Y.H., D.J, A.C., M.A., J.C., S.F., S.G., G.I., K.M., S.N., J-M.O., S.Sarkar, A.S., and N.T. performed sample isolation and processing. Q.X.X.L. pre-processed and analysed the first AIDA data freeze. E.V.B. and K.H.K. performed cell type annotation. R.S. performed data integration and SCENIC analyses. K.H.K. and M-S.P. performed eQTL analyses. D.R., S.Sankaran, and N.A.R. contributed towards study design. D.R. and N.A.R. developed and optimised protocols. Y.Tomofuji and Y.O. performed genotype QC and imputation and led the X-chromosome inactivation escape project. Y.Tomofuji, M.N., K.I., and Y.O. assembled GWAS summary statistics. L.M.T., J.M., J-E.P., and M.C. contributed towards scRNA-seq data analysis. Z.C. contributed towards DEG analysis. B.L., C.T., Y.Z., and Y.Tong led the splicing project. C.T.Y.T., A.M.T., Y.Y.H., and A.L. analysed the SLAS-2 flow cytometry data. T.P.N. and R.C.H. led the SLAS cohort. A-C.V. contributed towards cell type annotation. F.A.W., B.L., and H-H.W. contributed towards statistical methods. A.M., J-E.P. and M.C. contributed towards scRNA-seq experimental data generation. J.C.C., P.C., K.Y., B.S., M.P., M.L., and the SG10K_Health Consortium led AIDA cohorts and supervised sample collection. S.P., W-Y.P., J.W.S., P.P.M., V.C., P.M., and C-C.H. led and supervised scRNA-seq data generation. S.P., W-Y.P., and J.W.S. designed the study and supervised research. K.H.K. and S.P. wrote the manuscript, with input from all authors.

## Inclusion and Ethics

Our research included local researchers across all study sites in the 5 countries (India, Japan, Singapore, South Korea, Thailand) in every aspect of the research process, including study design, study implementation, data ownership, and authorship. Community engagement has been led by our authors and long-standing collaborators in each country with expertise in epidemiology, population genetics, and human disease studies.

## Data availability

The AIDA Data Freeze v2 gene-cell matrix (1,265,624 cells from 619 India, Japan, Singaporean Chinese, Singaporean Malay, Singaporean Indian, South Korea, and Thai Asian donors and 6 distinct Lonza commercial controls) and donor metadata will be available via the Chan Zuckerberg (CZ) CELLxGENE data portal (we will provide the link upon publication). The earlier AIDA Data Freeze v1 gene-cell matrix and visualisation is available via the CZ CELLxGENE data portal at https://cellxgene.cziscience.com/collections/ced320a1-29f3-47c1-a735-513c7084d508. The AIDA Data Freeze v2 cell annotation metadata will be available via the Cell Annotation Platform (CAP at https://celltype.info/; we will provide the link upon publication). The open-access AIDA datasets are available via the Human Cell Atlas Data Coordination Platform at https://data.humancellatlas.org/explore/projects/f0f89c14-7460-4bab-9d42-22228a91f185. The managed-access AIDA datasets are available via data access applications to the corresponding authors. The SLAS-2 dataset we analysed was profiled in a published study^43^, and is available through a data access application to the Gerontology Research Programme at the National University of Singapore, Singapore, and the Singapore Immunology Network (SIgN), A*STAR, Singapore.

## Code availability

The code for this manuscript is available at https://github.com/prabhakarlab/AIDA_Phase1.

## Methods

### Healthy donors in the Asian Immune Diversity Atlas

Living healthy human donors from India, Japan, Singapore, South Korea, and Thailand were profiled for this study. All study protocols were approved by the Institutional Review Boards (IRBs) of the institutions our laboratories are affiliated with (Genome Institute of Singapore: IRB 2020-012 and 2022-051; Nanyang Technological University: IRB-2016-11-030-01, IRB-2016-11-030, and 18IC4698; RIKEN: IRB H30-9; Samsung Genome Institute: IRB 2019-09-121; Faculty of Medicine Siriraj Hospital, Mahidol University: IRB 725/2563(IRB3); National Institute of Biomedical Genomics: IRB NIBMG/2022/1/0022) prior to dataset generation. Our sample collection sites were in Kolkata, Yokohama, Singapore, Seoul, and Bangkok, respectively. All donors provided written informed consent for sample and metadata collection and subsequent analyses. Donor metadata, such as age, female / male sex, self-reported ethnicity, height, weight, body mass index (BMI), and medication and dietary supplement consumption, were collected from donors via questionnaires and clinical measurements. We closely complied with all ethical regulations and our IRB conditions.

We included healthy donors in our atlas through applying the following exclusion criteria for donor datasets:

1) A person unable to provide informed consent.
2) A person with active infection or fever.
3) A person on regular medication (consumption of dietary supplements and / or herbal remedies was not considered in the exclusion of participants from our study).
4) A person with autoimmune disease.
5) A person with haemoglobin A1c (HbA1c)≥6%.

In addition, we excluded from our reference atlas persons who had received any vaccines in the 8 weeks prior to the date of blood draw.

We profiled 85 Singaporean Chinese, 70 Singaporean Indian, 61 Singaporean Malay, 149 Japan Japanese, 165 South Korea Korean, 59 Thailand Thai, and 30 India Indian donors for a total of 619 Asian donors for the AIDA Data Freeze v2 dataset.

We also included control PBMC samples from 6 distinct European donors (Lonza 4W-270, from lot numbers 3038099, 3038016, 3038097, 3038306, 3030004, and 3061635) in the AIDA Data Freeze v2 dataset.

### Isolation of peripheral blood mononuclear cells (PBMCs)

Eight ml of blood was drawn from each donor for this study using CPT tubes with sodium heparin (BD Vacutainer CPT, catalogue number 362753). Isolation of peripheral blood mononuclear cells (PBMCs) was performed according to a standardised protocol across all study sites. We used foetal bovine serum (FBS; Sigma-Aldrich catalogue number F2442) lot numbers 19G014 and 20A363 for both the PBMC isolation and the cell pooling and washing procedures. Briefly, blood samples collected in CPT tubes (standing in room temperature) were processed within 2 hours of blood collection using density-gradient centrifugation. Centrifugation steps were performed using a soft setting for centrifuge acceleration and deceleration. Blood samples were mixed 8-10 times prior to centrifugation at 1,500 x g in a horizontal rotor with swing-out head for 30 minutes at 20 °C. Plasma was aspirated; the PBMC layer was then collected, and spun down at 300 x g for 15 minutes at 20 °C. The cell pellet was resuspended in ACK lysing buffer (Thermo Fisher Scientific, catalogue number A10492) for lysis of red blood cells. Samples were washed twice with wash buffer (PBS pH 7.4, 1% FBS, 1 mM EDTA) and centrifuged at 300 x g for 15 minutes at 20 °C. PBMCs were cryopreserved in CryoStor CS10 cell freezing media (STEMCELL Technologies, catalogue number 079555). Cryovials were stored at −80 °C in controlled-rate cooling containers overnight before long-term storage in liquid nitrogen. We have made our detailed protocol harmonised across all study sites available via Protocols.io^87^ at https://www.protocols.io/view/pbmcs-isolation-from-cpt-tube-b8r9rv96.

### Single-cell experiments: Genetic multiplexing and sample pooling

Thawing and washing of individual PBMC samples and pooling (for genetic multiplexing) of donor samples were performed according to a standardised protocol across all study sites. Briefly, individual vials of PBMC donor samples were thawed in a 37 °C water bath for 1-2 minutes until no visible ice crystals were seen, and further thawed using pre-warmed thawing media (RPMI (Gibco catalogue number 21870076) + 5% Human Serum (Sigma-Aldrich catalogue number H4522) + 1% Penicillin-Streptomycin (Gibco catalogue number 15140122) + 1% L-Glutamine (Gibco catalogue number 25030081)). Individual samples were centrifuged at 300 x g for 5 minutes at 21 °C, and washed first with pre-warmed washing media (RPMI + 10% FBS + 1% Penicillin-Streptomycin + 1% L-Glutamine), followed by two washes with pre-warmed (PBS + 0.04% Bovine Serum Albumin (BSA, Capricorn Scientific catalogue number BSA-1S)). Individual samples were then each filtered through a 30 µm MACS SmartStrainer (Miltenyi Biotec) to remove cellular clumps and debris; samples were kept on ice for all the following procedures after filtering. Individual samples were counted with a 1:1 sample-to-trypan blue mix using an automated cell counter (Thermo Fisher Countess II FL), and resuspended to 1.50 × 10^6^ cells per ml in PBS + 0.04% BSA. Equal numbers and volumes of cells from each donor were pooled per experimental batch, and the pooled sample was counted using the same cell counting procedure as above prior to the 10x Genomics single-cell experiments. We have made our detailed protocol harmonised across all study sites available via Protocols.io^88^ at https://www.protocols.io/view/demuxlet-cell-preparation-protocol-b8sdrwa6.

### Single-cell experiments: 10x Genomics 5’ v2 RNA-sequencing, B-cell receptor sequencing (BCR-seq), and T-cell receptor sequencing (TCR-seq)

Fifteen Asian donors and one European control sample (Lonza 4W-270, from lot numbers 3038099, 3038016, 3038097, 3038306, 3030004, and 3061635; the first 5 lot numbers were used in AIDA Data Freeze v1, while all 6 lot numbers were used in AIDA Data Freeze v2) were pooled per batch, with two technical replicates (which we term as replicate *libraries*) performed for each batch. To allow for comparisons across donors from different population groups in the Singapore batches, we batch-randomised donors, ensuring that approximately the same numbers of Singaporean Chinese, Singaporean Malay, and Singaporean Indian donors (and the same age range and sex balance) were present per Singapore donor batch.

We loaded 40,000 cells from the donor PBMC pool for each technical replicate, and performed 10x Genomics 5’ v2 scRNA-seq, B-cell receptor sequencing (BCR-seq), and T-cell receptor sequencing (TCR-seq) experiments and library preparation according to the manufacturer’s protocols. We used a Chromium Controller at each study site for the 10x Genomics partitioning (generation of gel beads-in-emulsion (GEMs)) and barcoding procedures. We used the following 10x Genomics reagents for our experiments: Chromium Next GEM Chip K Single Cell Kit, Chromium Single Cell 5’ v2 Reagent Kit, Dual Index Kit TT Set A, Chromium Single Cell Human TCR Amplification Kit, and Chromium Single Cell Human BCR Amplification Kit. We quantified libraries using the Bioanalyzer 2100 with High Sensitivity DNA kits (Agilent Technologies). We pooled two 5’ v2 gene expression technical replicates (i.e., two libraries) per lane of an Illumina NovaSeq 6000 S4 flow cell. We pooled 20 BCR and / or TCR libraries per lane of an Illumina NovaSeq 6000 S4 flow cell. We sequenced the Japan, Singapore, South Korea, and Thailand libraries using a sequencing configuration of paired-end 150 bp with 10 bp dual i7 and i5 indices. Our pilot libraries from India were sequenced using a sequencing configuration of read 1: 26 bases and read 2: 90 bases with 10 bp dual i7 and i5 indices.

### Genomic DNA isolation, genotyping, genotype quality control, and genotype imputation

Genomic DNA was isolated from PBMCs from each donor using the QIAamp DNA Mini Kit (Qiagen, catalogue number 51306) according to the manufacturer’s protocol. Genotyping was performed using the Illumina GSAv3.0 array (Infinium Global Screening Array-24 Kit, catalogue number 20030770). We used the Illumina GenomeStudio version 2.0 software with the PLINK Input Report Plug-in v2.1.4 and the Infinium Global Screening Array v3.0 manifest files (BPM Format – GRCh38) to convert the raw data IDAT files to MAP and PED files. We used StrandScript^89^ to correct the Illumina genotyping data consistently to the GRCh38 human genome reference forward strand.

For our genetic demultiplexing workflow, we then used PLINK 1.9^90^ to retain autosomal variants present at a minimum minor allele frequency of 5% and to convert the data to VCF files. For each AIDA batch of donor samples, we included only SNPs with a 100% genotyping rate, and excluded indels. We corrected the reference allele base identity using bcftools norm -f^91^ with the GRCh38 reference genome, and removed any multi-allelic SNPs. We used the resulting VCF files as our input for genetic demultiplexing of individual single-cell sequencing libraries. We also used these VCF files for principal component analysis (PCA) of the AIDA genotype data, using the R prcomp function.

For genotype imputation, we performed both sample-level quality control (QC) and variant-level QC steps. Samples that had call ratios of below 0.98 (after considering autosomal variants with call ratios>0.99) were excluded from the imputation procedure. Related donor samples were identified through computation of the PI_HAT and Z1 identity-by-descent values using PLINK2^90^. During variant-level QC, variants with call ratio<0.99 were excluded. Variants that showed significant association with sex as well as variants with Hardy-Weinberg equilibrium (HWE) p-value<1e-6 were also excluded. Variants which had allele frequencies in the AIDA genotype dataset that differed from the 1000 Genomes hg38 dataset by more than 15% (for variants found in AIDA Singaporean Chinese, Japanese, and Korean donors versus the 1000 Genomes East Asian super-population, as well as variants found in AIDA Japanese donors versus the 1000 Genomes Japanese donors) or more than 17.5% (for variants found in AIDA Singaporean Indian donors versus the 1000 Genomes South Asian super-population) were excluded. In addition, variants for which we were unable to confidently match strand orientation to the 1000 Genomes hg38 dataset as well as duplicated variants were excluded from imputation. After these QC procedures, genotype imputation was performed using the Michigan Imputation Server^92^, utilising the 1000 Genomes hg38 (all populations) high-coverage reference panel (1000 Genomes Phase 3 (Version 5), with 2,504 samples and 49,143,605 sites on the autosomal chromosomes) as the imputation panel.

### Single-cell RNA-sequencing (scRNA-seq) pre-processing and quality control

We performed centralised pre-processing and quality control (QC) of all scRNA-seq datasets. We used the DRAGEN Single-Cell RNA pipeline in the Illumina DRAGEN v3.8.4 software (version 07.021.602.3.8.4-20-g74395e76) for pre-processing of paired-end FASTQ files from each individual scRNA-seq gene expression library from Japan, Singapore, South Korea, and Thailand to obtain one gene-cell matrix per library. We utilised the DRAGEN genetic demultiplexing workflow for detecting genetic doublets and for assigning cells to their donors based on the donor genotype data VCF file provided to the DRAGEN pipeline. We used GENCODE Release 32 (GRCh38, Ensembl 98, date 2019-09-05) as our gene annotation reference, and the associated GRCh38 primary genome assembly as our reference genome, and set --Aligner.hard-clips=0 and --Aligner.sec-aligns=3. We used the 737K-august-2016.txt barcode whitelist (corresponding to the list of barcodes relevant for the 10x Genomics Single Cell 5’ v2 assay) from the 10x Genomics Cell Ranger software installation in the DRAGEN pipeline. For the AIDA BCR-seq and TCR-seq Japan, Singapore, South Korea, and Thailand datasets, we processed the paired-end FASTQ files per library using Cell Ranger VDJ pipeline versions cellranger-5.0.0 and cellranger-5.0.1. For the AIDA scRNA-seq India datasets, as these could not be processed adequately through the DRAGEN pipeline, we ran Cell Ranger 7.0.1 with the default parameters, with introns included by default in the cellranger count step. For the AIDA BCR-seq and TCR-seq India datasets, we processed the FASTQ files using Cell Ranger VDJ pipeline version cellranger-4.0.0. We used the same Cell Ranger V(D)J reference (vdj_GRCh38_alts_ensembl-5.0.0) for all BCR and TCR datasets, and considered the high-confidence BCR and TCR contigs from the output files for our analyses.

For Japan, Singapore, South Korea, and Thailand single-cell experimental batches with all donor genotype data available, we used the DRAGEN genetic demultiplexing output for our genetic singlet and genetic doublet assignments. For batches with one missing donor genotype (e.g., due to problems with the genomic DNA extraction procedure), we used Freemuxlet^93^ (https://github.com/statgen/popscle) on the BAM files from the DRAGEN pipeline output, with the default Freemuxlet parameters, to assign cells to donors. We then performed genotype concordance analyses by comparing the Freemuxlet-inferred genotypes against the Illumina GSAv3 genotyping array data to match the Freemuxlet clusters to donors. For these Freemuxlet analyses, we used as our input VCF file into the dsc-pileup step the set of exonic variants that were present at a minor allele frequency ≥5% in the East Asian and / or the South Asian super-populations in the 20181203_biallelic_SNV GRCh38 version of the 1000 Genomes dataset (EBI European Variation Archive accession PRJEB30460). For the AIDA India datasets, we ran Demuxlet on the BAM output files from Cell Ranger with the default parameters, except for setting --group-list as the list of barcodes (barcodes.tsv.gz) from the unfiltered Cell Ranger output. We excluded any library with excessively high genetic doublet rates from all downstream analyses.

We performed QC of our scRNA-seq dataset in two stages. We first performed library-level QC by analysing each individual library. We filtered out cells for which fewer than 300 GENCODE Release 32 genes were detected (number of detected genes (NODG)<300). We identified a preliminary cell type annotation for each library for use in our doublet identification workflow. For AIDA Data Freeze v1, we identified the top 2,000 highly variable features using the variance-stabilising transformation option in the Seurat 4.1.1 R package^94^, scaled the data using all genes, and then performed principal component analysis on these highly variable features. We performed nearest-neighbour analyses based on the resulting principal components, and ran Louvain clustering in Seurat at a resolution of 1.0. For AIDA Data Freeze v2, to guard against batch-to-batch variability in heterotypic doublet identification, we instead performed reference projection of the scRNA-seq library to a reference panel of immune cell transcriptomes using the RCAv2 software^95^, performed nearest-neighbour analyses based on the principal components of the reference projection coefficients, and ran Louvain clustering in Seurat at a resolution of 1.0. For both data freezes, we annotated the resulting clusters based on a majority vote of the major cell type annotation labels assigned by RCAv2 to cells within each cluster.

We used the genetic doublet proportion for a library (combining the proportions of mixed genetic identity and ambiguous identity droplets) to estimate the likely total doublet rate for that library (proportion of genetic doublets in the library out of all cells with NODG≥300 multiplied by (number of donor samples)/(number of donor samples-1))^96^. We used this estimate of total doublets in a library, as well as the RCAv2 reference projection-based clustering and annotation of clusters (for estimation of homotypic doublet proportion) as our input into DoubletFinder, which we used for identifying heterotypic doublets^97^. We then removed cells that had more than 10 (*HBA1* UMIs + *HBB* UMIs), since these cells could be red blood cells, or cells contaminated with red blood cell RNA transcripts. We checked for any sample swaps by examining the number of singlets per donor (typically ∼1,000 singlets and almost always >>100 singlets per donor per library, even if a particular donor sample had low cell viability after thawing). We also examined for concordance of the scRNA-seq-inferred female / male sex (inferred from the ratio of total counts from non-pseudoautosomal region (PAR) Y-chromosome genes to total counts of PAR Y-chromosome genes) with the donor metadata. By checking for matches between the transcriptomic and genotype data, our integrative analysis of scRNA-seq reads with genotyping array output through the genetic demultiplexing workflow helped guard against any inter-batch sample swaps.

After we performed the library-level QC procedures, we performed cell type-specific QC on our dataset. We removed any cell that was flagged as a doublet by the DRAGEN genetic demultiplexing workflow or by the DoubletFinder workflow from our downstream analyses, and included only single cells from healthy donors that had provided written informed consent and had not withdrawn consent from the study. We then combined single cells from multiple libraries across countries, performed reference projection of such combinations of cells to a reference panel of immune cell transcriptomes using the RCAv2 software^95^, and performed nearest-neighbour analyses based on the principal components of the reference projection coefficients. We ran Louvain clustering in Seurat^94^ at a resolution dependent on the size of the combination of cells, increasing the resolution for a larger set containing more cells. We annotated the resulting clusters based on a majority vote of the major cell type annotation labels assigned by RCAv2 to cells within each cluster. We performed cell type-specific QC on all single cells across all libraries by applying per-cell NODG and percentage mitochondrial read (pMito) filters that were manually determined for each major cell type (B, CD34^+^ haematopoietic stem and progenitor cell (HSPC), myeloid (including both monocyte and conventional dendritic cell), natural killer (NK), plasma cell, plasmacytoid dendritic cell (pDC), platelet, T); these major cell types were largely distinct in gene expression space (except for NK and T cells) in our scRNA-seq analyses. For example, our NODG filters excluded any myeloid cell with NODG<500, and any other leukocyte with NODG<1,000. Our pMito filters excluded any cell with pMito>12.5% (for plasma cells and platelets) and pMito>8% (for all other major cell types). We included only cells within our designated NODG and pMito range for cell type annotation and downstream analyses.

Following these two stages of QC, AIDA Data Freeze v1 comprised 1,058,909 PBMCs from 503 Asian donors and 5 European controls profiled in Japan, Singapore, and South Korea, which we made available to the research community pre-publication (<u>Supplementary Note</u>), including via the CZ CELLxGENE data portal and as part of the first CZ CELLxGENE Census^25^. AIDA Data Freeze v2, which we focused on in this study, comprises 1,265,624 PBMCs from 619 Asian donors and 6 European controls (median number of detected genes (NODG) per library: 1342-2296, median NODG of 93 libraries=2003; median percentage mitochondrial reads (pMito) per library: 2.07%-4.08%, median pMito of 93 libraries=3.53%) (**Figure S1A**). We had a median of 122 high-confidence BCR barcodes and 986 high-confidence TCR barcodes per donor (**Figure S1B**).

### Cell population-specific quality control, data integration, sub-clustering, and cell type annotation

We performed cell population-specific quality control (QC), feature selection, data integration, sub-clustering, and annotation on the following cell populations separately: 1) B cells; 2) pDCs and myeloid cells; and 3) ILC, NK, and T cells. For the ILC, NK, and T cells, we performed a second round of data integration as well as re-clustering on the following two cell populations separately: 3a) CD4^+^ T and dnT cells, and 3b) ILC, NK, and T cells that were neither CD4^+^ T nor dnT cells.

We utilised genes that were expressed by ≥0.1% of cells in our cell populations of interest for our analyses. We first identified genes expressed by ≥0.1% of cells in each of the following cell types: B, pDC, myeloid, NK, and T. We retained the union of these genes for the combined pDC and myeloid cell populations, and also the combined NK and T cell populations. We then re-normalised the respective gene-cell matrices (B; pDC and myeloid; NK and T) by the total expression of retained genes.

We excluded cells with heightened platelet marker gene expression by performing a platelet marker gene QC step in the first round of the sub-clustering analyses. We separately identified platelet gene expression distributions in 1) B cells, 2) pDCs and myeloid cells, and 3) ILC, NK, and T cells. We excluded cells that were amongst the top 30% of cells with a non-zero sum of expression of four platelet marker genes (*ITGA2B*, *PF4*, *PPBP*, *TUBB1*) from the first round of cell population-specific data integration, sub-clustering, and cell type annotation procedures.

Following the platelet marker gene QC step, we re-identified the genes that were expressed by ≥0.1% of cells in our cell populations of interest. This was performed prior to each data integration procedure.

We performed data integration using the Seurat anchor integration reciprocal principal component analysis (RPCA) algorithm^28^. We integrated across all scRNA-seq libraries and treated each scRNA-seq library as a batch.

For each library, we performed log-normalisation and identification of the top 2000 highly variable features using the variance-stabilising transformation option in Seurat. We selected integration features using the Seurat SelectIntegrationFeatures function, scaled each library and ran PCA on each library using these integration features, and selected the library with the highest number of cells for the cell population of interest as our reference dataset. We identified integration anchors via Seurat RPCA using this reference dataset and the first 30 principal components. We then performed data integration using Seurat IntegrateData and its default parameters (e.g., the first 30 dimensions as well as k.weight=100 for the number of neighbours considered in the anchor weighting procedure). For control analyses, we performed Harmony for data integration^44^, treating each scRNA-seq library as a batch. We chose highly variable genes (HVG) by their prevalence across the highly variable genes for all scRNA-seq libraries (Harmony-HVG), followed by Harmony integration, and sub-clustering and cell type annotation in the Harmony-HVG embedding.

We performed sub-clustering using the integrated embedding principal components, and identified marker genes for one cluster versus all other clusters using the implementation of the single-cell Wilcoxon rank-sum test in the Seurat FindMarkers differential gene expression function^94^. We then annotated our cells based on marker genes curated from the literature as well as through examination of gene expression across clusters in our dataset (**<u>Table S2</u>**). Our annotation framework involved four hierarchical levels for annotating sub-clusters (**Figures S2A,B**). At the most detailed level (Level 4), we named and described individual sub-clusters, which we refer to as cluster identities in this study. We manually merged these towards cell subtypes widely-recognised in the literature at higher hierarchical levels in the PBMC clustering hierarchy. We used well-defined cell type descriptors from the literature (e.g., naïve, memory) as well as marker genes and their heightened / lowered levels of expression (hi / lo, respectively) for nomenclature. We considered the fold-change in expression of marker genes for a cluster, as well as the proportion of cells within a cluster that expressed the marker genes of interest, based on single-cell Wilcoxon rank-sum tests, for our annotation nomenclature (e.g., log_2_(fold-change)>0.5 and >50% of a cluster expressing a gene for a “hi” annotation of that gene). For distinguishing T cell clusters, we considered the proportion of cells within a cluster that had high-confidence TCR barcodes. We annotated clusters in detail, where possible, but left annotations at a coarser level or described subtypes as “unknown” (e.g., CD4+_T_unknown instead of a specific CD4^+^ T subtype) when there was no compelling evidence in support of a more detailed annotation. We also flagged clusters that appeared to have 1) heightened platelet gene expression; 2) the combination of (a) a low range of NODG values, (b) low expression of canonical marker genes that should be expressed in the cell type, and (c) heightened expression of genes that, relative to other clusters, tend to be highly expressed in PBMC scRNA-seq datasets (e.g., the long non-coding RNA genes *MALAT1*, *NEAT1*); as well as 3) clusters with heightened expression of marker genes from other lineages or cell types (e.g., T cells that showed heightened expression of monocyte marker genes) for caution in downstream interpretation.

### Cell type proportion analyses

For our analyses, we focused on population groups with at least 50 donors, donors with at least 800 cells passing our quality control filters, and cell subtypes with at least ∼10 cells per donor on average. We utilised linear models of self-reported ethnicity, age, female / male sex, and their two-way interaction terms to examine the correlation of these dimensions of human diversity with the log_10_(Proportion) of immune cell types and subtypes. We evaluated several denominators, including all PBMCs per donor as well as all NK, T, and ILC cells (for NK and T cell subtypes) to guard against possible changes in overall myeloid or lymphoid cell abundance across population groups.

To compute the variance in cell subtype proportions explained by a dimension of human diversity, we examined the multiple R-squared values (equivalent to the variance explained) from linear regression models in which only a single dimension (e.g., age, self-reported ethnicity, BMI, or sex) was considered. We focused on Singapore donors for this analysis, using total PBMCs without platelets as our denominator. For validation of the impact of self-reported ethnicity on cell subtype proportions, we analysed the Singapore Longitudinal Ageing Study wave-2 (SLAS-2)^98^ flow cytometry dataset profiled in a published study^43^. This dataset included 824 donors spanning 55-94-years, comprising 719 SG_Chinese, 40 SG_Indian, and 65 SG_Malay donors. In the aforementioned study, briefly, flow cytometry data was analysed using the FlowJo software (BD). Cell populations were gated in FlowJo and the event counts of each cell population were exported into Microsoft Excel for calculating frequencies of cell populations. We used a model of log_10_(Proportion)∼Age+Sex+Individual_Self_reported_ethnicity (e.g., SG_Chinese or SG_Indian or SG_Malay), and the PBMCs/Single_Cells/Live_Cells/CD34^+^CD45^+^ event count (total leukocytes) as our denominator for these analyses. This corresponded to the same linear model featuring total PBMCs without platelets as our denominator in the AIDA scRNA-seq analyses of the Singapore population groups.

Statistical tests, including computation of two-tailed t-test and two-tailed Wilcoxon rank-sum test p-values, were performed in R. Plots were generated using the ggplot2 R package^99^ and the R plotting functions.

### Cell neighbourhood enrichment analyses

We performed two types of cell neighbourhood enrichment analyses. We first examined the impact of single dimensions of human diversity across the whole AIDA atlas. We performed Seurat RPCA integration for all cells and all libraries according to our Seurat RPCA workflow described in the *Cell population-specific quality control, data integration, sub-clustering, and cell type annotation* Methods section. From the resulting integrated embedding, we considered only population groups with at least 50 donors for the cell neighbourhood enrichment analysis. We identified the 500 nearest neighbours for each cell in our atlas using the first 30 principal components from our integrated embedding, and computed the number of cells corresponding to our dimension of human diversity of interest (i.e., female or male sex, one of the self-reported ethnicities, one of the four age ranges for our dataset (in years, ages 19 to 32, 33 to 40, 41 to 49, 50 to 77) and the number of cells corresponding to the complement (i.e., all other cells from the other sex considered, the other self-reported ethnicities, or the other age ranges, respectively). We normalised the ratio of these two values against the ratio of total cells in our atlas corresponding to the dimension of human diversity of interest, to the total number of cells corresponding to the complement. We then performed a log_2_ transformation of these values, and overlaid these values on gene expression UMAPs.

Next, we used MiloR version 1.5^46^ to test for differential cell neighbourhood abundance using models accounting for multiple dimensions of human diversity. The MiloR analysis allowed for overlapping cell neighbourhoods and computation of spatial false-discovery rates (FDR). We implemented MiloR for all combinations of cell populations (1) B; 2) pDC and myeloid; 3) CD4^+^ T and dnT; and 4) ILC, NK, and T cells that were neither CD4^+^ T nor dnT cells) with dimensions of human diversity. We performed the MiloR analyses using the Seurat RPCA integrated embedding from the cell type annotation workflow, following the removal of cells with heightened platelet gene expression and data integration using the Seurat RPCA anchor integration procedure. We performed the following workflow for donors with at least 800 cells per donor. We set k=900, such that the peak in the histogram of neighbourhood sizes was ∼3,000, which was a number ∼5 times the number of donors analysed through the MiloR workflow. We used a model of Age+Sex+Self_reported_ethnicity for differential cell neighbourhood abundance testing, with SG_Chinese, SG_Indian, or SG_Malay substituted into the self-reported ethnicity term when investigating cell neighbourhood enrichment associated with a particular self-reported ethnicity. We treated age as a continuous variable when examining the effects of female / male sex or self-reported ethnicity, and categorised age into 50-77 years versus <50 years when examining the effects of age. We ran MiloR with the graph-based sampling refinement scheme for identifying neighbourhoods, and with the graph-overlap option for the spatial FDR weighting scheme. We identified the most enriched and least enriched neighbourhoods for each combination of cell population and dimension of human diversity of interest, paying particular attention to neighbourhoods with spatial FDR<0.1. We also examined these enrichment patterns in the context of our sub-clustering annotations for generating beeswarm plots. We performed differential gene expression analyses for identifying neighbourhood-associated marker genes using the single-cell Wilcoxon rank-sum test implemented in Seurat FindMarkers, comparing the cells within the neighbourhood of interest (annotated via a majority vote of the cell subtype annotation of cells within the neighbourhood), against all other cells that had been annotated as being of that same cell subtype. We visualised cell neighbourhood enrichment for each population group by plotting the log_2_ transformation of the mean fold-change for each cell across all the overlapping neighbourhoods to which that cell belonged.

To examine the impact of the Singapore self-reported ethnicity covariates, we performed the above MiloR workflow on the Singapore donors (with at least 800 cells per donor) only.

### Differential gene expression analyses

We performed edgeR (R package version 3.38.4)^54^ analyses of pseudobulk gene expression data; the edgeR log-likelihood pseudobulk testing workflow was assessed in a benchmarking study to show good performance in terms of reducing the number of false discoveries^100^. We considered only Singapore donors with at least 800 PBMCs passing quality control for our edgeR analyses, to minimise confounding with technical variation across study sites. For each cell subtype, we considered only donors with at least 10 cells for the cell subtype of interest. We then obtained pseudobulk profiles by aggregating gene-cell count matrices into gene-donor count matrices for each cell type or subtype of interest. We pre-filtered our gene list to remove lowly-expressed genes: we retained genes that were expressed in at least 10% of donors per cell subtype after pseudobulk aggregation, and further filtered out any genes that had fewer UMIs than the total number of donors considered for differential gene expression analysis for a cell subtype.

We incorporated age, sex, and scRNA-seq experimental batch in the edgeR generalised linear model, and tested for one Singapore self-reported ethnicity population group (SG_Chinese, SG_Malay, or SG_Indian) compared to the other two Singapore groups to analyse differential gene expression across population groups. We computed false-discovery rates (FDR) by performing Benjamini-Hochberg correction^57^ of edgeR p-values per cell type or subtype.

We used clusterProfiler version 4.4.4^101^ for performing gene-set enrichment analyses (GSEA) and visualisation of GSEA results. We used the fgseaMultilevel option with the Gene Ontology (GO) Biological Processes terms and Benjamini-Hochberg-corrected false-discovery rates^57^ for our GSEA analyses. For each self-reported ethnicity-cell subtype combination, we supplied a pre-ranked gene list, comprising all genes tested per combination for differential gene expression, ranked by the -log_10_(edgeR p-value) multiplied by the sign of the edgeR-log_2_ fold-change value.

### Differential transcription factor activity analyses

We implemented a SCENIC-based^62^ workflow to investigate differential transcription factor activity based on our scRNA-seq gene expression data, using the AIDA Data Freeze v1 dataset for these analyses. We used the pySCENIC version of SCENIC^102^, starting with a curated list of 1,390 transcription factors that was a subset of a list of human transcription factors^103^, and utilising the Motif2TF v10 annotations and the hg38 refseq_r80 SCENIC+ motif databases with a search space of 10 kb flanking the transcription start sites of genes. We performed GRNBoost2 and cisTarget using default parameters to prioritise transcription factors and their target genes of interest (collectively known as a “regulon” per transcription factor). We examined the regulon activity per cell type of interest using AUCell, and compared the distributions of the median of the raw AUCell scores per cell subtype per donor across different population groups in a cell type-specific manner via two-tailed Wilcoxon rank-sum tests performed in Python. We computed Benjamini-Hochberg-corrected false-discovery rates^57^ for all tests performed for a regulon of interest.

For our SCENIC workflow, we assessed several parameters to check the robustness of our inferences of regulons that we identified as being of interest. We ran a minimum of 10 trials of GRNBoost2 in determining the regulon sets of target genes. We varied the proportion of the genes in the gene-ranking of each cell considered by the AUCell computation procedure, examining the output from considering the top 5%, 10%, and 15% of genes in the gene-ranking. We identified GRNBoost2 regulons both using a combined gene-cell matrix spanning all cells from all Singapore libraries, as well as subsets of cells from each Singapore population group. We tested regulons from multiple trials of GRNBoost2, as well as the union of regulon target genes across trials. We also tested both a pseudobulk input (summing, within the gene-cell matrix, across all cells of a particular cell subtype per donor to obtain a gene-donor matrix for each cell subtype) as well as the original scRNA-seq input for our AUCell computations. In this manuscript, we report findings that have been observed in multiple SCENIC GRNBoost2 trial-AUCell analysis combinations.

### Single-cell pseudobulk expression quantitative trait loci (eQTL) pipeline

We developed a single-cell pseudobulk expression quantitative trait loci (eQTL) pipeline, using the AIDA Data Freeze v1 dataset for these analyses. For the gene expression values, we computed pseudobulk values per cell subtype of interest. We first filtered out genes that were expressed by <1% of cells in the cell subtype of interest, as well as donors and cells for which there were fewer than 10 cells per donor of the cell subtype of interest. We normalised each remaining gene by the total number of UMIs of retained genes per cell, and computed a mean gene expression value per donor from the cells of the subtype of interest. We performed log1p transformation with a scale factor of 10,000 to approximate a normal distribution of mean gene expression values for each donor. For the genetic variants of interest, after the genotype data quality control and imputation procedures described above, we retained autosomal, biallelic variants that had minor allele frequencies ≥5% in the AIDA cohort. We removed related donors (with this being computed per cell subtype after the above-mentioned donor cell number filters were applied).

We used Matrix eQTL (version 2.3 R package)^104^ for association testing and FDR computation to identify cell type-specific cis-eQTLs within 1 Mb of the gene of interest, retaining the results from all tests performed, regardless of p-value, for downstream analyses. We used the Matrix eQTL additive linear model, which returned t-statistics and two-tailed t-test p-values. Our eQTL model featured the following as covariates: age, sex, self-reported ethnicity and / or country, top 10 genotype PCs, and top 10 gene expression PCs (computed per cell subtype). We also performed eigenMT^68^ to prioritise one lead SNP per gene, using the authors’ default parameters.

We performed eQTL replication by comparing our AIDA eQTLs against eQTLs identified in the DICE^69^ and ImmuNexUT^70^ projects. The DICE eQTLs had been uniformly processed as part of the EMBL-EBI eQTL Catalogue^105^. We compared our AIDA naïve B eQTLs against eQTL Catalogue dataset QTD000474, AIDA CD4^+^ T naïve eQTLs against eQTL Catalogue dataset QTD000479, AIDA CD8^+^ T naïve eQTLs against eQTL Catalogue dataset QTD000489, AIDA CD14^+^ Monocyte eQTLs against eQTL Catalogue dataset QTD000504, and AIDA CD16^+^ NK eQTLs against eQTL Catalogue dataset QTD000509. For eQTL replication using the ImmuNexUT datasets, we analysed NBDC Human Database dataset E-GEAD-420. For AIDA eQTLs that had a FDR<0.05 within the cell subtype of interest, we identified data from DICE / ImmuNexUT with a SNP ID-gene combination match, and compared the beta values from the AIDA dataset against the corresponding DICE / ImmuNexUT dataset. For the GTEx v8 whole blood eQTL^18^ comparison, we analysed eQTL Catalogue dataset QTD000356. Since we were comparing eQTLs from all 20 AIDA cell subtype datasets against the GTEx v8 whole blood eQTL dataset, we applied a more stringent criteria for selection of AIDA eQTLs for comparison, restricting our analysis to the set of AIDA eQTLs with FDR<0.05 (Benjamini-Hochberg-adjusted^57^ p-values) across all tests in all cell subtypes.

We obtained the 1000 Genomes Project super-population allele frequencies from ENSEMBL release 105 (https://ftp.ensembl.org/pub/release-105/variation/vcf/homo_sapiens/, file date 20210906), converted these alternate allele frequencies to minor allele frequencies by taking the lower of the two allele frequencies per reference-alternate pair, and computed allele frequencies for any AIDA eQTL for which we were able to identify a SNP ID match in the 1000 Genomes VCF file.

### Colocalisation of eQTL with variants identified through genome-wide association studies (GWAS)

We compiled summary statistics from genome-wide association studies (GWAS) of immune-related diseases (asthma^77^, rheumatoid arthritis^74^, systemic lupus erythematosus (SLE)^106^, Graves’ disease^80^, atopic dermatitis and type 1 diabetes^107^) versus controls, and hospitalised COVID-19 cases versus the general population^108^, all of which included an Asian cohort within a trans-ancestry study, or were performed entirely by studying Asian donors (**<u>Table S7</u>**). We used coloc version 5.2.3, utilising the approximate Bayes factor^109^ enumeration implementation of the coloc R package that assumes at most a single causal variant per trait^110^. We performed colocalisation analyses of GWAS and eQTL (each eGene) traits. We used coloc to compute the posterior probabilities of each possible scenario of significant genetic association with traits in the eQTL analysis and in the GWAS analysis. We paid particular attention to the posterior probability (PP) for the case where both the gene expression (eQTL) and disease risk (GWAS) traits are associated with and share a single causal variant, which we abbreviated as “colocalisation PP”. We used LocusCompareR^111^ to visualise colocalisation events, setting the population parameter to “EAS” and selecting the hg38 option, given the format and scope of our GWAS summary statistics.

### Context-dependent eQTL analyses

We used the AIDA Data Freeze v1 dataset for these analyses. We examined significant eQTL-eGene pairs (FDR<0.05 per cell subtype), identified through our Single-cell pseudobulk expression quantitative trait loci (eQTL) pipeline, for context-dependent eQTL effects. We intersected eQTL-eGene pairs from naïve B, *IGHM*^hi^ memory B, and *IGHM*^lo^ memory B, analysing a total of 54,798 SNP-gene pairs. We performed the following smoothing and gene-gene correlation procedures for memory B cells from each country (Japan, Singapore, and South Korea) in AIDA Data Freeze v1 separately to minimise the influence of batch effects on the assembly of gene modules. We conducted k-nearest neighbour-based smoothing of the gene-cell matrix, calculating smoothed expression values for each cell by averaging the gene expression of each cell and its nearest 30 neighbours. We performed gene-gene correlation analyses, and then averaged the gene-gene correlation matrices across the three countries to obtain a single gene-gene correlation matrix. We thereafter used WGCNA^112^ to identify gene modules, which we used to model cellular contexts and cell states.

To identify context-dependent eQTL effects, we obtained module scores per memory B cell by averaging the expression values of genes belonging to the module of interest. We tested for the interaction between donor genotype and module score on gene expression per cell using univariate Poisson models^72^, implemented via lme4::glmer^113^ in R:

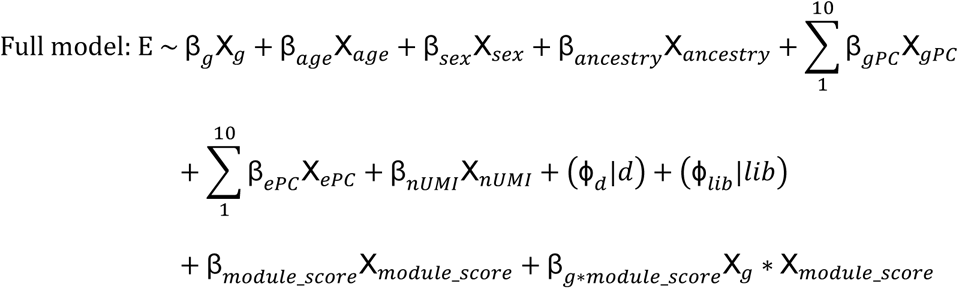

E represents each gene of interest’s UMI counts in the cell of interest, *g* represents donor genotype, *gPC* represents genotype PC, *ePC* represents gene expression PC based on the PCA of the gene-cell matrix, *nUMI* represents the cell’s total number of UMI counts, *d* represents donor, and *lib* represents scRNA-seq library. All covariates are modelled as fixed effects, except for donor and library, which are modelled as random effects.

The null model was computed using the same covariates as the full model, leaving out only the β_*g*∗*module*_*score*_X_*g*_ ∗ X_*module*_*score*_ term. P-values were calculated for each full model-null model pair using the anova function in R. We computed Benjamini-Hochberg-corrected false-discovery rates^57^ across all p-values.

## Quantification and statistical analysis

All statistical tests were performed using R, R packages, or Python, with the specific testing details listed in the individual method sections. All statistical tests performed were two-tailed.

## Supplementary Figures and Figure Legends

**Supplementary Figure S1:**
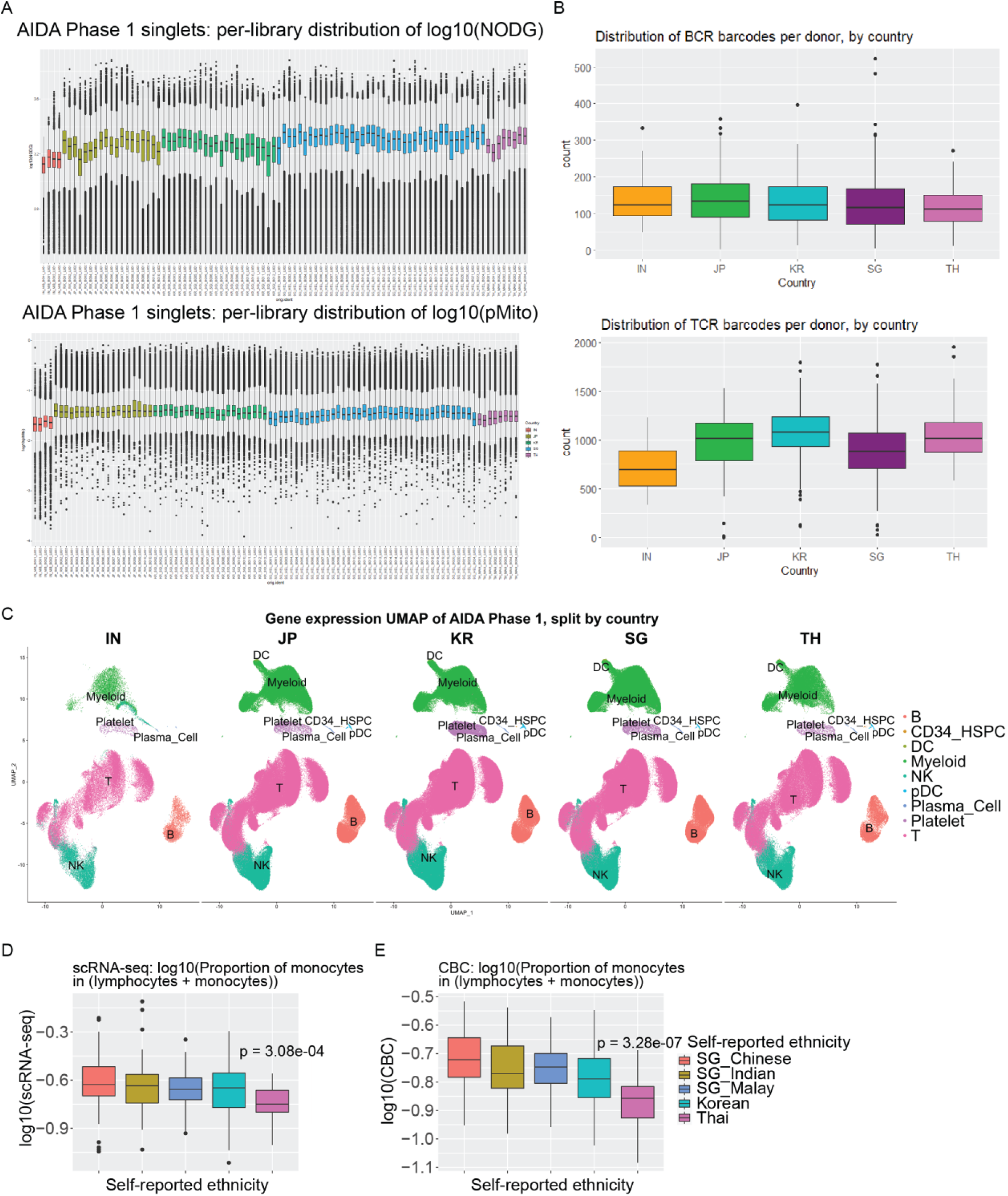
An scRNA-seq reference atlas of circulating immune cells from healthy Asian donors. (**A**) Distributions of (top) log_10_(numbers of detected genes (NODG) per cell) and (bottom) log_10_(percentage mitochondrial UMIs out of all UMIs per cell (pMito)) in AIDA Data Freeze v2 libraries. (**B**) Distributions of high-confidence BCR (top) and TCR (bottom) barcodes per donor in each country. (**C**) The same UMAP as that in Figure 2A, split by country where donors were profiled. Distributions of log_10_(proportion of monocytes out of all lymphocytes plus monocytes per donor) in (**D**) AIDA scRNA-seq and (**E**) complete blood count (CBC) data, categorised by donor self-reported ethnicity. P-values indicate results from two-tailed Wilcoxon rank-sum tests. Boxplots depict the median via the thickest centre horizontal line, the first and third quartiles as the bottom and top of the box respectively, and 1.5x the interquartile range through the whiskers; outliers are depicted as single points.

**Supplementary Figure S2:**
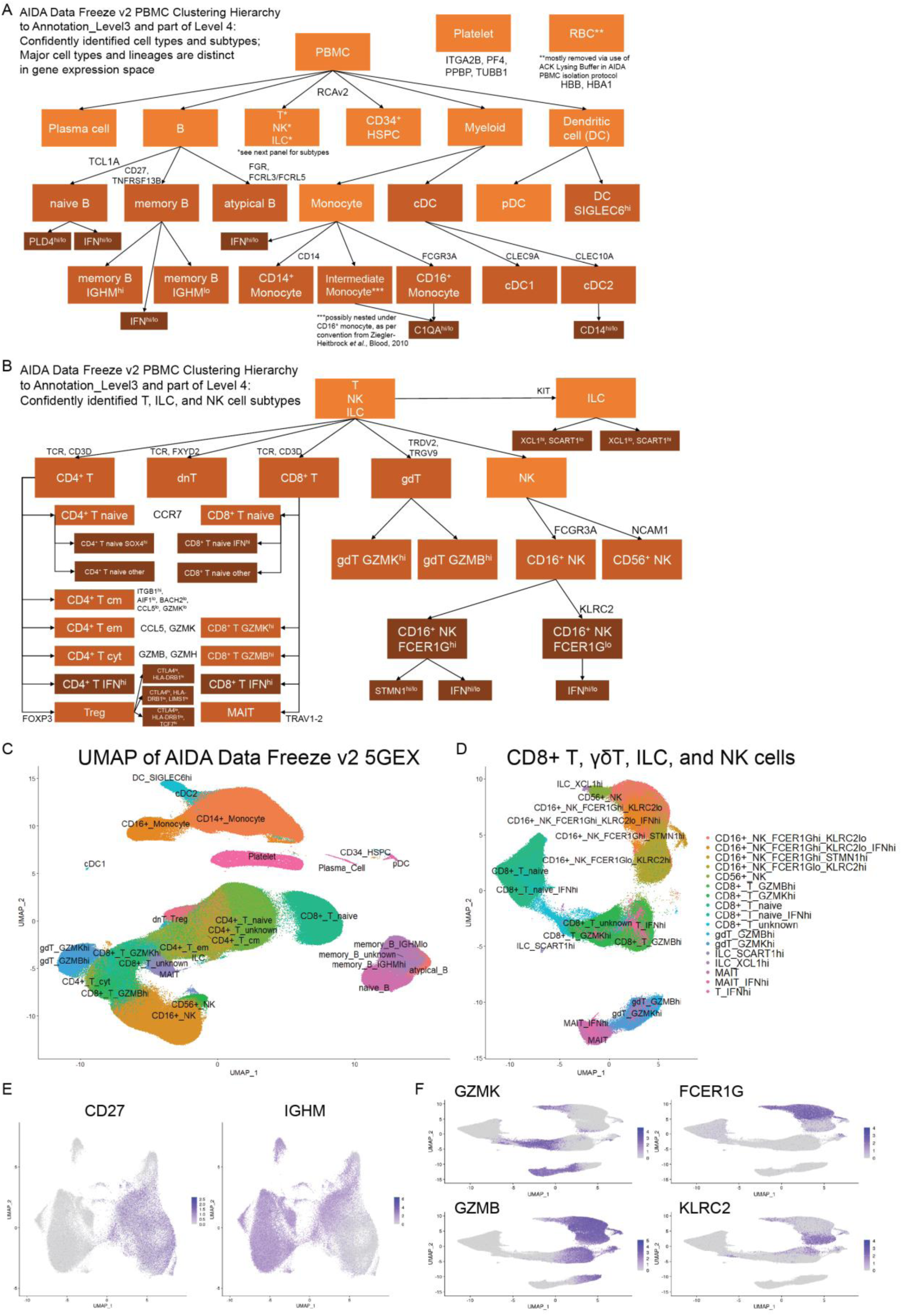
AIDA cell type annotation metadata. PBMC cell type clustering hierarchy in AIDA, with major marker genes indicated, for (**A**) all PBMCs and (**B**) ILC, NK, and T cell subtypes. Boxes coloured in orange indicate cell types, boxes coloured in light brown indicate cell subtypes, and boxes coloured in dark brown indicate more granular cluster identities. (**C**) Gene expression UMAP of the AIDA dataset, labelled by AIDA Level 3 cell type annotations. (**D**) UMAP of CD8^+^ T, γδT, ILC, and NK cells, labelled by AIDA Level 4 cell type annotations. Feature plots of (**E**) *CD27* and *IGHM*, overlaid on UMAPs of B cells, and (**F**) features representing immune cell gradients (*GZMK*, *GZMB*, *FCER1G*, and *KLRC2*), overlaid on UMAPs of CD8^+^ T, γδT, ILC, and NK cells; intensities of colours correspond to the log-normalised gene counts per cell.

**Supplementary Figure S3:**
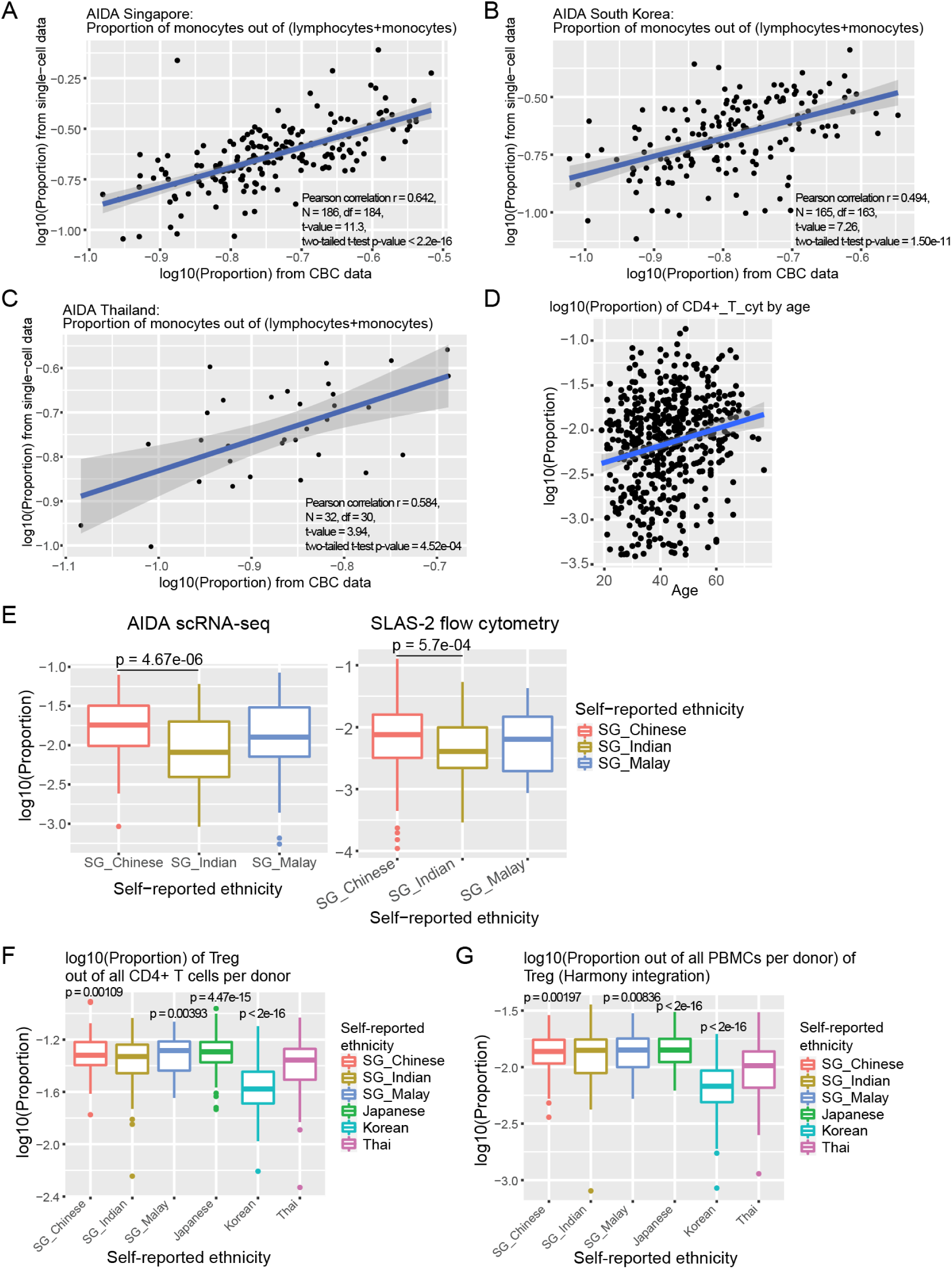
Impact of human diversity on cell subtype proportions. Scatterplots of proportion of monocytes (out of all monocytes and lymphocytes) per donor in the scRNA-seq datasets (y-axis) versus that in matched complete blood counts (x-axis) for AIDA donors in (**A**) Singapore, (**B**) South Korea, and (**C**) Thailand. (**D**) Scatterplot of CD4^+^ T cytotoxic (CD4+_T_cyt) cell proportions against donor age for all AIDA donors. (**E**) Boxplots depicting MAIT cell proportions across Singapore self-reported ethnicities in (left) our AIDA scRNA-seq dataset and (right) the SLAS-2 flow cytometry dataset; two-tailed t-test p-values adjacent to lines indicate comparisons of two population groups. (**F**) Boxplots depicting the proportions of regulatory T (Treg) cells out of all CD4^+^ T cells per donor across all population groups. (**G**) Boxplots depicting the proportions of Treg cells out of all PBMCs per donor across all population groups, after data integration using Harmony^44^ and re-clustering and re-annotation of cells. Scatterplots are overlaid with blue linear regression lines; grey bands indicate the 95% confidence intervals. Boxplots depict the median via the thickest centre horizontal line, the first and third quartiles as the bottom and top of the box respectively, and 1.5x the interquartile range through the whiskers; outliers are depicted as single points. Two-tailed t-test p-values in (**F**,**G**) are for the self-reported ethnicity covariate in a model of log10(Proportion)∼Age+Sex+Individual_Self_reported_ethnicity.

**Supplementary Figure S4:**
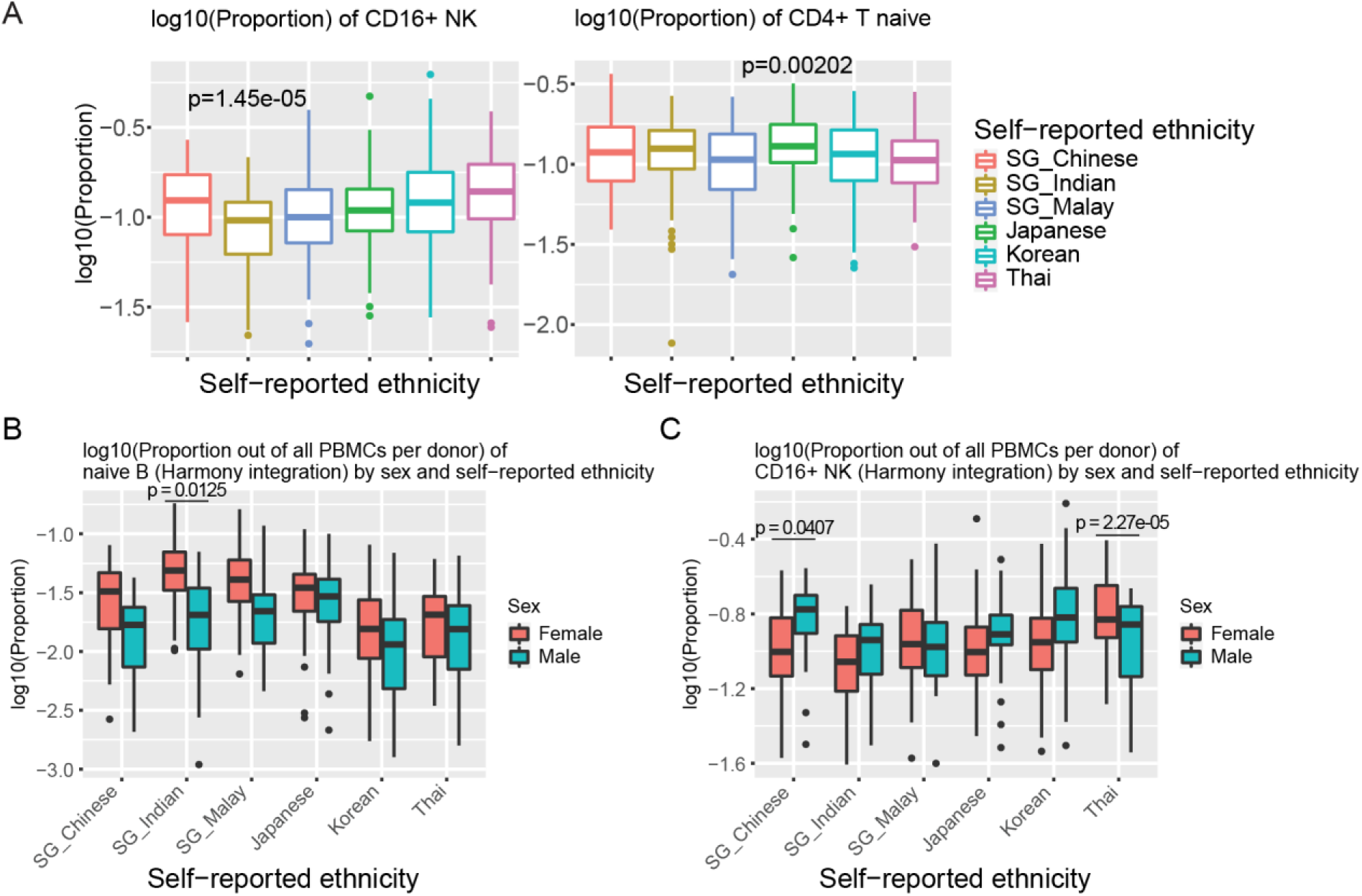
Impact of self-reported ethnicity and sex on cell subtype proportions. (**A**) Boxplots depicting (left) CD16^+^ NK and (right) CD4^+^ T naïve proportions out of all PBMCs per donor across self-reported ethnicities. Boxplots depicting (**B**) naïve B and (**C**) CD16^+^ NK proportions out of all PBMCs per donor across all population groups and female / male sex, after data integration using Harmony^44^ and re-clustering and re-annotation of cells. Boxplots depict the median via the thickest centre horizontal line, the first and third quartiles as the bottom and top of the box respectively, and 1.5x the interquartile range through the whiskers; outliers are depicted as single points. Two-tailed t-test p-values in (**A**) pertain to the self-reported ethnicity covariate in a model of log_10_(Proportion)∼Age+Sex+Individual_Self_reported_ethnicity. Two-tailed t-test p-values adjacent to lines pertain to the interaction terms between sex and individual population groups in (**B**,**C**).

**Supplementary Figure S5:**
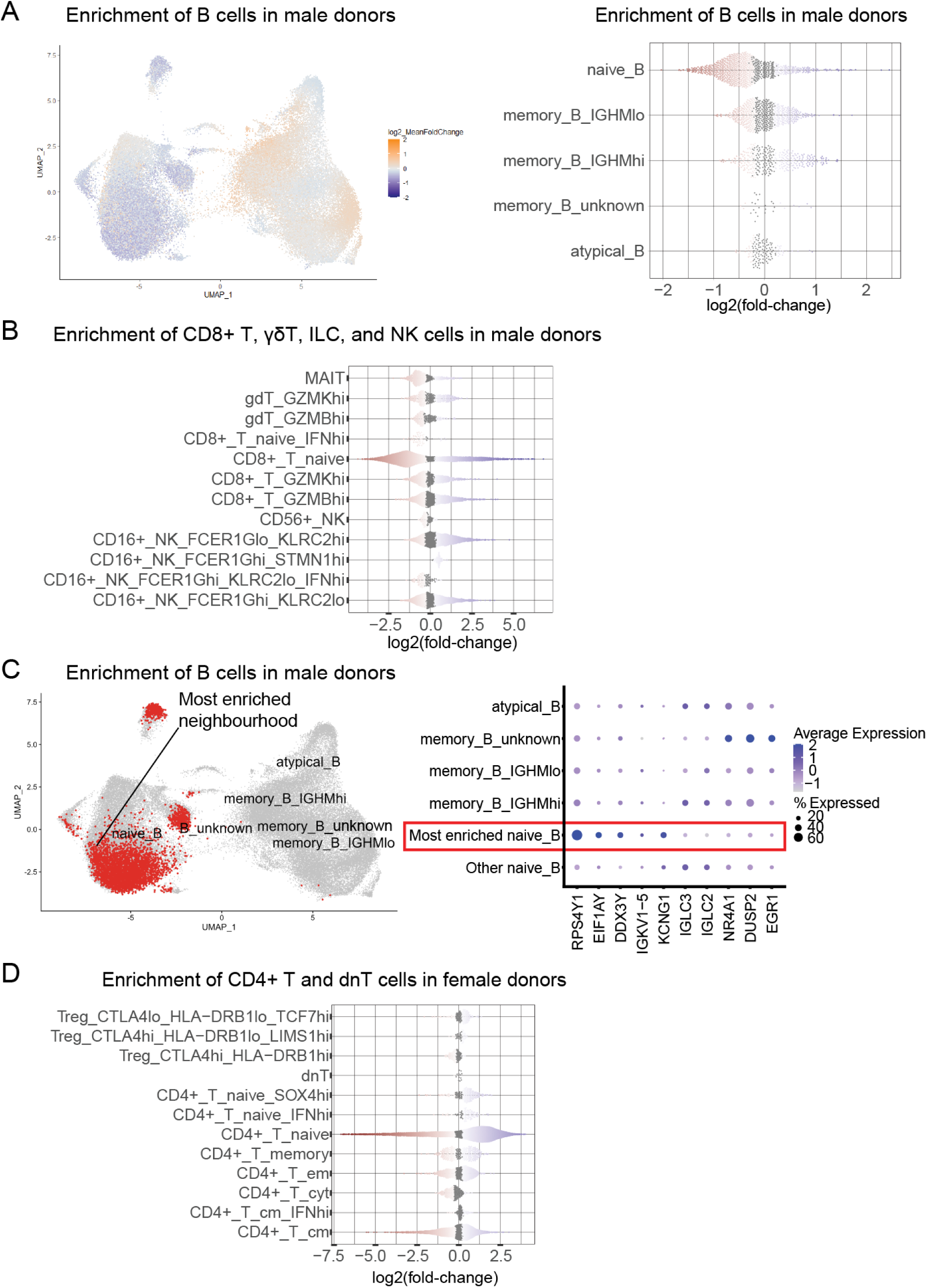
Single-cell signatures of sex. (**A**) (Left) Gene expression UMAP and (right) beeswarm plot depicting enrichment of B cell neighbourhoods in males versus females. (**B**) Beeswarm plot depicting enrichment of CD8^+^ T, γδT, ILC, and NK cell neighbourhoods in males versus females. (**C**) (Left) Cells highlighted within a UMAP, corresponding to the most enriched B cell neighbourhood in males versus females. (Right) Dot plot of top 5 upregulated and top 5 downregulated genes (as compared to all other naïve B cells) of the most enriched B cell neighbourhood in males versus females. (**D**) Beeswarm plot depicting enrichment of CD4^+^ T and dnT cell neighbourhoods in females versus males. For the UMAP depicting cell neighbourhood enrichment, each cell is coloured by their log_2_(mean fold-change) value for all overlapping cell neighbourhoods that the cell was grouped in. Orange hues indicate cell neighbourhood enrichment for the population group, while blue hues indicate cell neighbourhood depletion; darker hues correspond to higher magnitudes of enrichment or depletion, capped at log_2_(mean fold-change)=|2|. For beeswarm plots, each point corresponds to one cell neighbourhood; cell neighbourhoods are classified by the majority cell type annotation within the neighbourhood. Points coloured in red (depletion of neighbourhood for the dimension of human diversity of interest) and in blue (enrichment of neighbourhood) correspond to spatial FDR values<0.1.

**Supplementary Figure S6:**
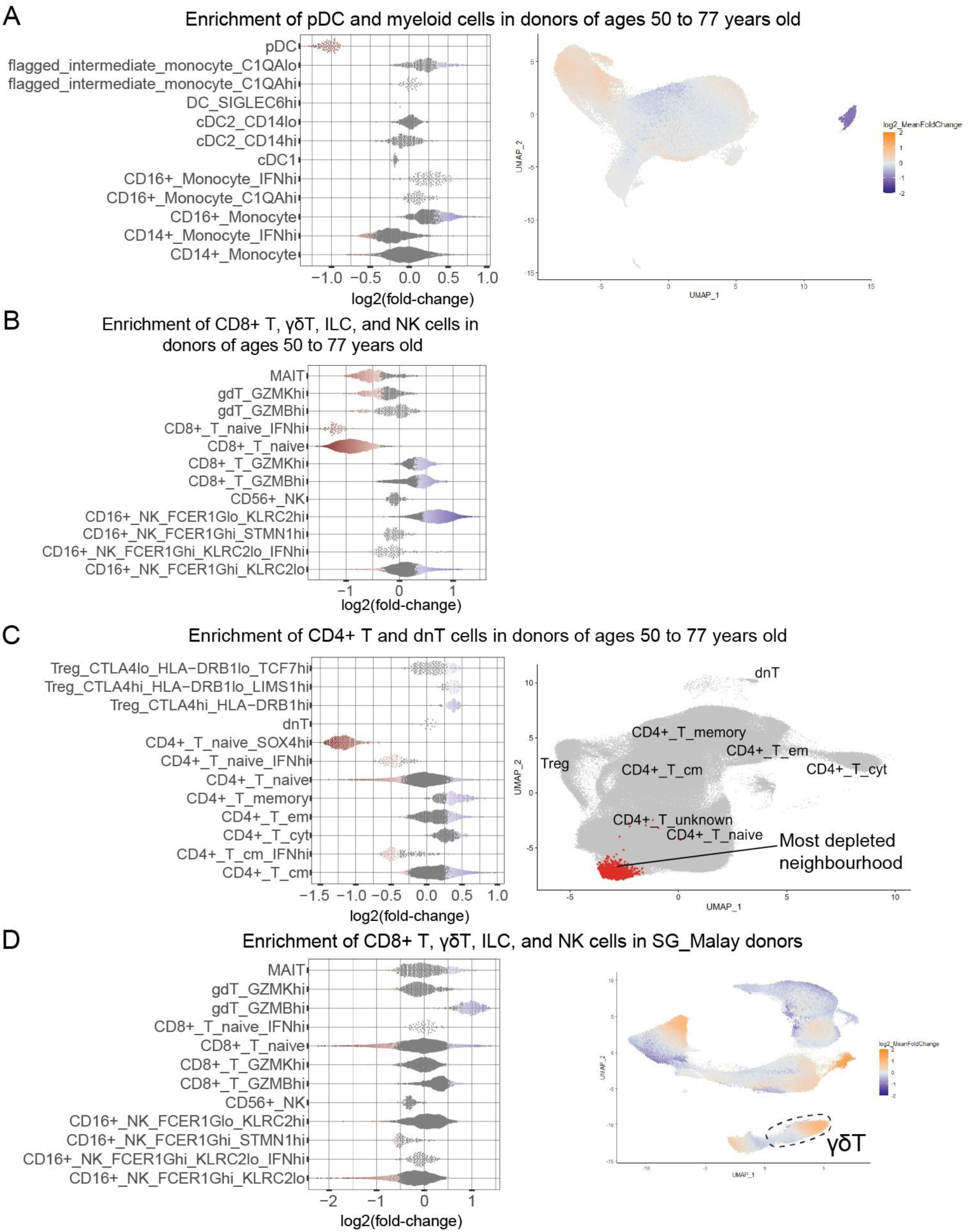
Single-cell signatures of age and self-reported ethnicity. (**A**) (Left) Beeswarm plot and (right) gene expression UMAP depicting enrichment of pDC and myeloid cell neighbourhoods in donors ≥50-years-old versus younger donors. Beeswarm plots depicting enrichment of (**B**) CD8^+^ T, γδT, ILC, and NK, and (**C**) (left) CD4^+^ T and dnT cell neighbourhoods in donors ≥50-years-old versus younger donors. (**C**) (Right) Cells highlighted within a UMAP, corresponding to the most depleted CD4^+^ T and dnT cell neighbourhood in donors ≥50-years-old versus younger donors. (**D**) (Left) Beeswarm plot and (right) UMAP depicting enrichment of CD8^+^ T, γδT, ILC, and NK cell neighbourhoods in SG_Malay donors, based on analysis of all AIDA donors; γδT cells are indicated by dashed lines. For UMAPs depicting cell neighbourhood enrichment, each cell is coloured by their log_2_(mean fold-change) value for all overlapping cell neighbourhoods that the cell was grouped in. Orange hues indicate cell neighbourhood enrichment for the population group, while blue hues indicate cell neighbourhood depletion; darker hues correspond to higher magnitudes of enrichment or depletion, capped at log_2_(mean fold-change)=|2|. For beeswarm plots, each point corresponds to one cell neighbourhood; cell neighbourhoods are classified by the majority cell type annotation within the neighbourhood. Points coloured in red (depletion of neighbourhood for the dimension of human diversity of interest) and in blue (enrichment of neighbourhood) correspond to spatial FDR values<0.1.

**Supplementary Figure S7:**
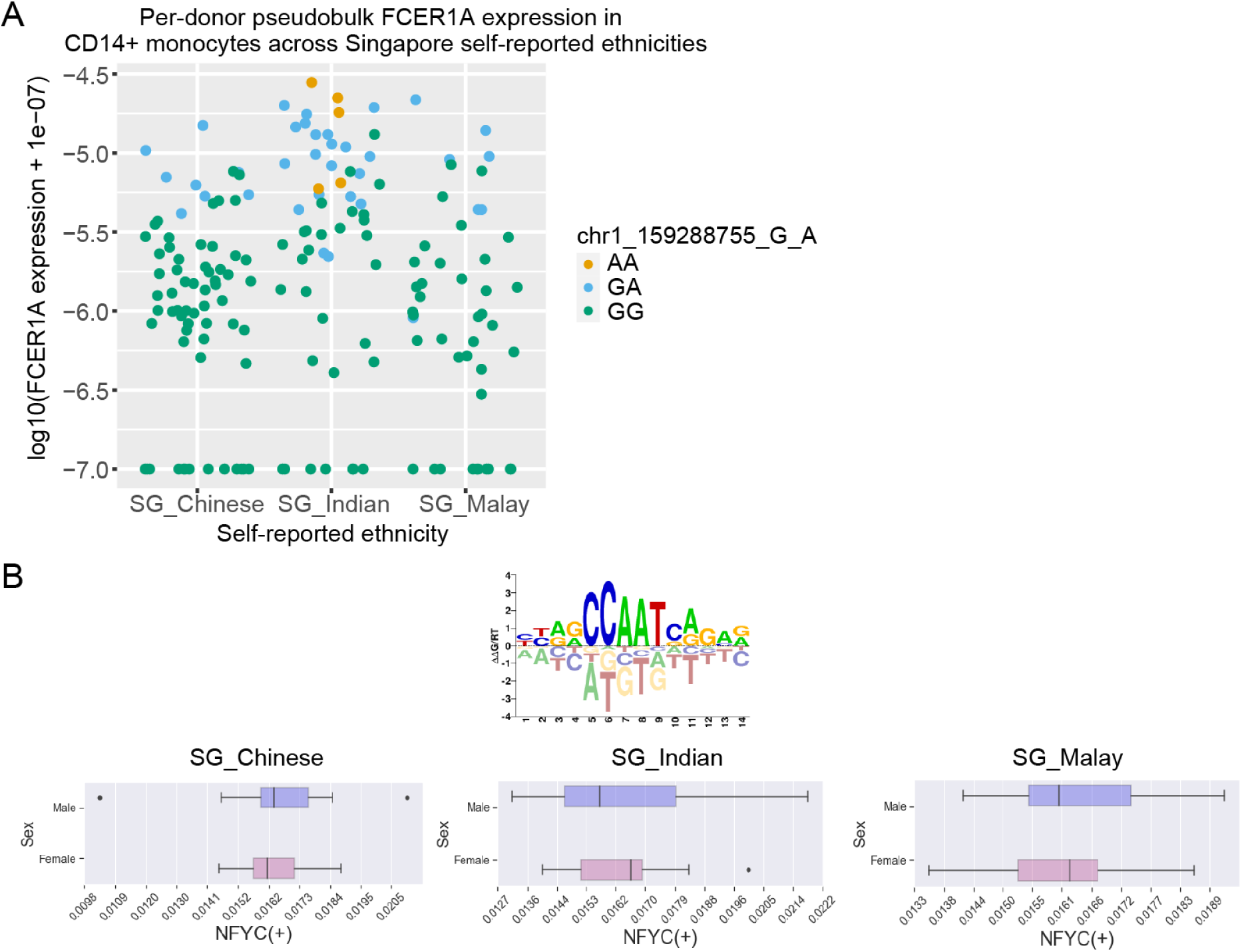
Cell type-specific molecular variation across population groups. (**A**) Jitter plots of log_10_ transformation of per-donor pseudobulk normalised *FCER1A* expression values (with added pseudocount of 1e-07) in CD14^+^ monocytes across the Singapore self-reported ethnicities, for donors with imputed genotype data available. Each dot represents a donor; dots are coloured by the donor genotype for the chr1_159288755_G_A locus. (**B**) (Top) NFYC transcription factor binding site motif from CIS-BP^58^ (M09442_2.00), and (bottom left to bottom right) boxplots depicting the median *NFYC* AUCell score across all regulatory T (Treg) cells per donor for each of the indicated Singapore population groups. Boxplots depict the median via the thickest centre vertical line, the first and third quartiles as the left side and right side of the box respectively, and 1.5x the interquartile range through the whiskers; outliers are depicted as single points. These boxplots depict the output from one SCENIC GRNBoost2 trial-AUCell analysis combination.

**Supplementary Figure S8:**
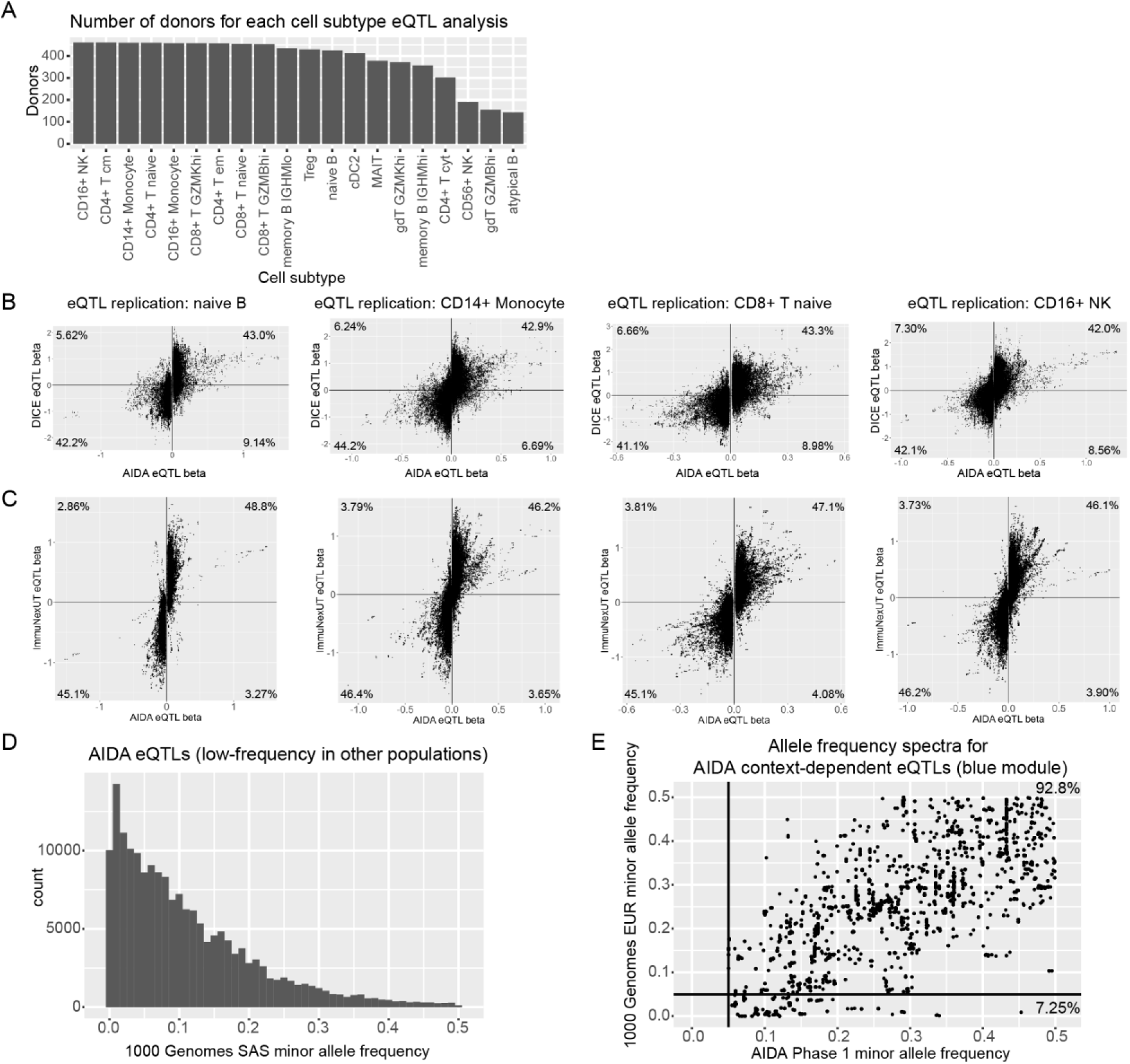
Replication of AIDA pseudobulk eQTL analyses and identification of population-specific eQTLs. (**A**) Bar charts of the number of donors analysed for AIDA eQTLs per cell subtype. Scatterplots of (**B**) DICE^69^ (y-axis) and (**C**) ImmuNexUT^70^ (y-axis) versus AIDA (x-axis) eQTL effect size (beta) values of SNP-gene pairs with AIDA eQTL FDR<0.05 per cell subtype, for (left to right) naïve B, CD14^+^ monocyte, CD8^+^ T naïve, and CD16^+^ NK. Percentages of all SNP-gene pairs that lie within a quadrant are indicated. (**D**) Histogram of minor allele frequencies (maf) in the 1000 Genomes South Asian (SAS) super-population, for AIDA eQTLs that were low frequency (maf 1%-5%) or rare (maf<1%) in at least one of the 1000 Genomes African, Admixed American, or European (EUR) super-populations. (**E**) Scatterplot depicting allele frequency spectra (1000 Genomes EUR super-population maf on y-axis, and AIDA Phase 1 cohort maf on x-axis) for AIDA context-dependent eQTLs (modulated by blue module score). Percentages of eQTLs with 1000 Genomes EUR maf≥0.05 and maf<0.05 are indicated. Bold lines indicate maf=0.05.

## Supplementary Tables

<u>Supplementary Table S1:</u> AIDA donor metadata: donor DCP_ID, self-reported ethnicity, age, country (study site), female / male sex, BMI, and scRNA-seq experimental batch.

<u>Supplementary Table S2:</u> Marker genes used for cell type annotation in AIDA.

<u>Supplementary Table S3:</u> Comparison of AIDA scRNA-seq and SLAS-2 flow cytometry cell types.

<u>Supplementary Table S4:</u> A diversity atlas reference: the relationships of self-reported ethnicity (controlling for age and sex) with circulating immune cell subtype proportions.

<u>Supplementary Table S5:</u> List of Singapore self-reported ethnicity-associated differentially expressed genes identified through edgeR analyses of pseudobulk values per cell type per donor (FDR<0.05 per cell subtype).

<u>Supplementary Table S6:</u> List of 143,918 SNP-gene pairs that have FDR<0.05 from Benjamini-Hochberg correction of eigenMT-corrected p-values.

<u>Supplementary Table S7:</u> List of curated disease GWAS summary statistics incorporated in colocalisation analyses.

<u>Supplementary Table S8:</u> Colocalisation analyses of AIDA eQTL and GWAS: posterior probability >80% of both traits being associated with and sharing a single causal variant.

<u>Supplementary Table S9:</u> List of SG10K_Health consortia authors.

## Supplementary Note

We released the first AIDA data freeze (“AIDA Data Freeze v1 dataset”) to the community pre-publication, via the CZ CELLxGENE data portal as well as the Human Cell Atlas Data Portal. The AIDA Data Freeze v1 dataset was also part of the first CZ CELLxGENE Census assembled in May 2023. We profiled 75 Singaporean Chinese, 60 Singaporean Indian, 54 Singaporean Malay, 149 Japan Japanese, and 165 South Korea Korean donors for a total of 503 Asian donors for the AIDA Data Freeze v1 dataset.

Going from AIDA Data Freeze v1 to AIDA Data Freeze v2, we excluded 5 Asian donors from v1 (SG_HEL_H141, SG_HEL_H185, SG_HEL_H203, SG_HEL_H239, and SG_HEL_H347) with ambiguous medication data. We added 121 new Asian donors (32 Singapore donors, 59 Thai donors, and 30 India Indian donors). These new Asian donors included donors SG_HEL_H262 and SG_HEL_H269, as well as donors profiled in experimental batches SG_HEL_B023, SG_HEL_B024, TH_MAH_B001, TH_MAH_B002, TH_MAH_B003, TH_MAH_B004, IN_NIB_B001, and IN_NIB_B002. We also removed two libraries with high doublet rates (SG_HEL_B011_L002 and SG_HEL_B021_L001).

The AIDA Data Freeze v1 gene-cell matrix (1,058,909 cells from 503 Japan, Singaporean Chinese, Singaporean Malay, Singaporean Indian, and South Korea Asian donors and 5 distinct Lonza commercial controls), with BCR-seq and TCR-seq metadata, and donor age, sex, and self-reported ethnicity metadata, is available via the Chan Zuckerberg (CZ) CELLxGENE data portal at https://cellxgene.cziscience.com/collections/ced320a1-29f3-47c1-a735-513c7084d508.

